# Distinct SAP102 and PSD-95 nano-organization defines multiple types of synaptic scaffold protein domains at single synapses

**DOI:** 10.1101/2023.09.12.557372

**Authors:** Sarah R. Metzbower, Poorna A. Dharmasri, Aaron D. Levy, Michael C. Anderson, Thomas A. Blanpied

## Abstract

The MAGUK family of scaffold proteins plays a central role in maintaining and modulating synaptic signaling, providing a framework to retain and position receptors, signaling molecules, and other synaptic components. Of these scaffold proteins, SAP102 and PSD-95 are essential for synaptic function at distinct developmental timepoints and perform overlapping as well as unique roles. While their similar structures allow for common binding partners, SAP102 is expressed earlier in synapse development and is required for synaptogenesis, whereas PSD-95 expression peaks later in development and is associated with synapse maturation. PSD-95 and other key synaptic proteins organize into subsynaptic nanodomains that have a significant impact on synaptic transmission, but the nanoscale organization of SAP102 is unknown. How SAP102 is organized within the synapse, and how it relates spatially to PSD-95 on a nanometer scale, could impact how SAP102 clusters synaptic proteins and underlie its ability to perform its unique functions. Here we used DNA-PAINT super-resolution microscopy to measure SAP102 nano-organization and its spatial relationship to PSD-95 at individual synapses. We found that like PSD-95, SAP102 accumulates in high-density subsynaptic nanoclusters. However, SAP102 nanoclusters were smaller and denser than PSD-95 nanoclusters across development. Additionally, only a subset of SAP102 nanoclusters co-organized with PSD-95, revealing that within individual synapses there are nanodomains that contain either one or both proteins. This organization into both shared and distinct subsynaptic nanodomains may underlie the ability of SAP102 and PSD-95 to perform both common and unique synaptic functions.

**Significance statement:** SAP102 and PSD-95 are two key members of the MAGUK family of synaptic scaffold proteins that are critical for synapse development, maintenance, and modification during plasticity. Because PSD-95 has a highly complex subsynaptic nanostructure that impacts synaptic function, we asked if SAP102 is similarly organized into nanoclusters at individual synapses and how it relates to PSD-95 within synapses. We found that SAP102 forms subsynaptic nanoclusters with unique properties, and that within individual synapses proteins both concentrate into overlapping nanodomains, as well as form MAGUK-specific nanodomains. This demonstrates that organization of synaptic proteins into nanoclusters is likely to be maintained within the family of MAGUK proteins and reveals potential mechanism for specializing functions within individual synapses based on scaffold protein nanodomains.

## Introduction

The membrane-associated guanylate kinase (MAGUK) synaptic scaffold proteins are among the most abundant synaptic proteins and establish a structural foundation on which the rest of the synapse is built (Sheng & Hoogenraad, 2007; Won et al., 2017). The family includes PSD-95, PSD-93, SAP102, and SAP97, which act as essential mediators of synaptic development, function, receptor recruitment, and plasticity by performing both overlapping and unique roles at the synapse (Carlisle et al., 2008; Chen et al., 2021; Cuthbert et al., 2007; Elias et al., 2008; Levy et al., 2015).

These proteins are distributed throughout the postsynaptic density (PSD) (Chen et al., 2008; Zhang & Diamond, 2009) and interlink with other scaffold families to establish the PSD as a flexible matrix in which transmembrane proteins are positioned (Blanpied et al., 2008; Kerr & Blanpied, 2012; Won et al., 2017). A prominent feature of PSD-95 is its presence in high-density assemblies termed nanoclusters (NCs) (Broadhead et al., 2016; M. Fukata et al., 2015; MacGillavry et al., 2013; Nair et al., 2013). PSD-95 NCs accumulate with other postsynaptic proteins to anchor glutamate receptors (Dai et al., 2019; Fukata et al., 2013; MacGillavry et al., 2013; Nair et al., 2013) and align them with vesicle release-related presynaptic proteins (Tang et al., 2016) to promote glutamate exposure (Ramsey et al., 2021). Despite this clear role for PSD- 95 nanostructure in synaptic function, the nano-organization of other MAGUK family members is unknown.

PSD-95 and SAP102 exemplify the complex functional relationship among MAGUKs. These structurally similar scaffolds are frequently at the same synapse (Cizeron et al., 2020) and interact with a highly overlapping set of synaptic proteins (Elias et al., 2008; Lau & Zukin, 2007; Su et al., 2018; J. Zhu et al., 2016). However, they are each critical at distinct developmental timepoints (Cizeron et al., 2020; Elias et al., 2006; Petralia et al., 2005; Sheng & Hoogenraad, 2007), as SAP102 recruits AMPARs and NMDARs during synaptogenesis, while PSD-95 performs this role at established synapses (Elias et al., 2008). SAP102 plays an additional role in synaptic NMDAR removal that PSD-95 does not (Chen et al., 2012), and while both PSD-95 and SAP102 can support LTP, there is evidence they do so via distinct mechanisms (Chen et al., 2021; Cuthbert et al., 2007). Moreover, specific multimolecular complexes formed by each scaffold have been suggested to underlie brain postsynaptic diversity (Zhu et al., 2018).

Understanding the basis for the functional divergence of SAP102 and PSD-95 will require knowing whether their nano-organization within the PSD is similar. While the proteins are structurally homologous, there are differences that could impact their subsynaptic organization. PSD-95 palmitoylation at an N-terminal di-cysteine motif is required for its synaptic and subsynaptic organization (Balderas et al., 2022; Fukata et al., 2015) whereas SAP102 is not palmitoylated (El-Husseini et al., 2000). Both interact with NMDARs, but SAP102 shows a preference for GluN2B-containing NMDARs (GluN2B-NMDARs) (Sans et al., 2000), while PSD- 95 tends to interact with GluN2A-containing NMDARs (GluN2A-NMDARs) (Elias et al., 2008; Gardoni & Di Luca, 2021). NMDAR subtypes have been reported to occupy distinct subsynaptic domains (Kellermayer et al., 2018), thus suggesting that SAP102 and PSD-95 may also occupy distinct nanodomains. Work with STED and electron microscopy indicated that PSD-95 and SAP102 are enriched at different distances vertically from the synaptic membrane (Zheng et al., 2010, 2011), though their lateral organization and relationship to one another have not been investigated.

Here we used DNA-PAINT super-resolution microscopy to directly visualize the nanoscale distribution and relationship of SAP102 and PSD-95 at individual synapses in cultured hippocampal neurons. We found that SAP102, like PSD-95, is organized into high-density subsynaptic NCs. However, SAP102 NCs have distinct properties and do not consistently co-localize with PSD-95 NCs. While both proteins show nanostructural changes across development, their spatial relationship to each other was largely consistent. This organization supports a model in which some SAP102 NCs can carry out synaptic functions distinct from PSD-95, while within common nanodomains, PSD-95 and SAP102 may play overlapping roles due to their mutual proximity to common interacting proteins.

## Materials and Methods

### Primary Neuron Culture

All animal procedures were approved by the University of Maryland Animal Use and Care committee. Dissociated hippocampal neuron cultures were prepared from both male and female E18 Sprague Dawley rat embryos (Charles River) as described previously (Dharmasri et al., 2023; Divakaruni et al., 2018). Neurons were plated on glass coverslips (18 mm #1.5, Warner Instruments) coated with poly-l-lysine at 30,000 cells per coverslip. Cells were grown in Neurobasal A + GlutaMax, gentamycin, and B27 supplement, with FUDR added at 1 week to suppress glial growth. Cells were fixed between 13-15 DIV for most experiments, between 6-8 or 20-22 DIV for developmental comparisons, and at 21 DIV for experiments with bassoon labeling.

### Reagents and antibodies

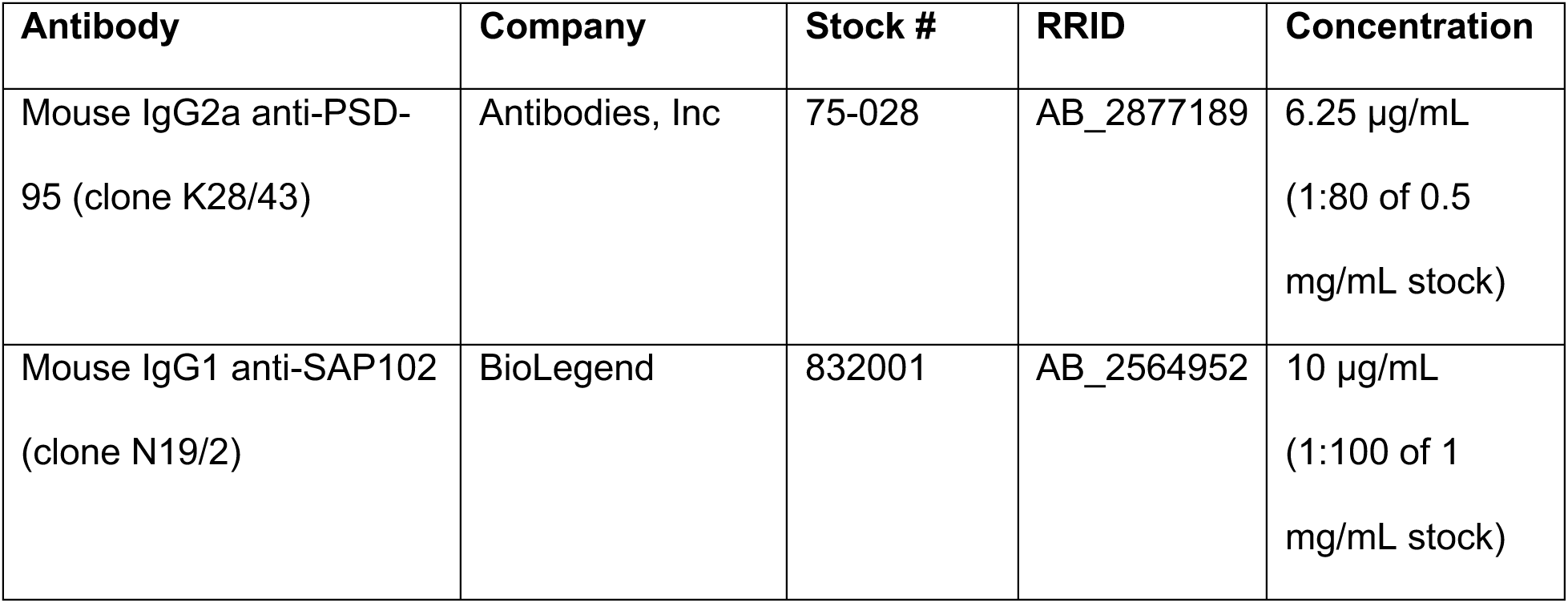
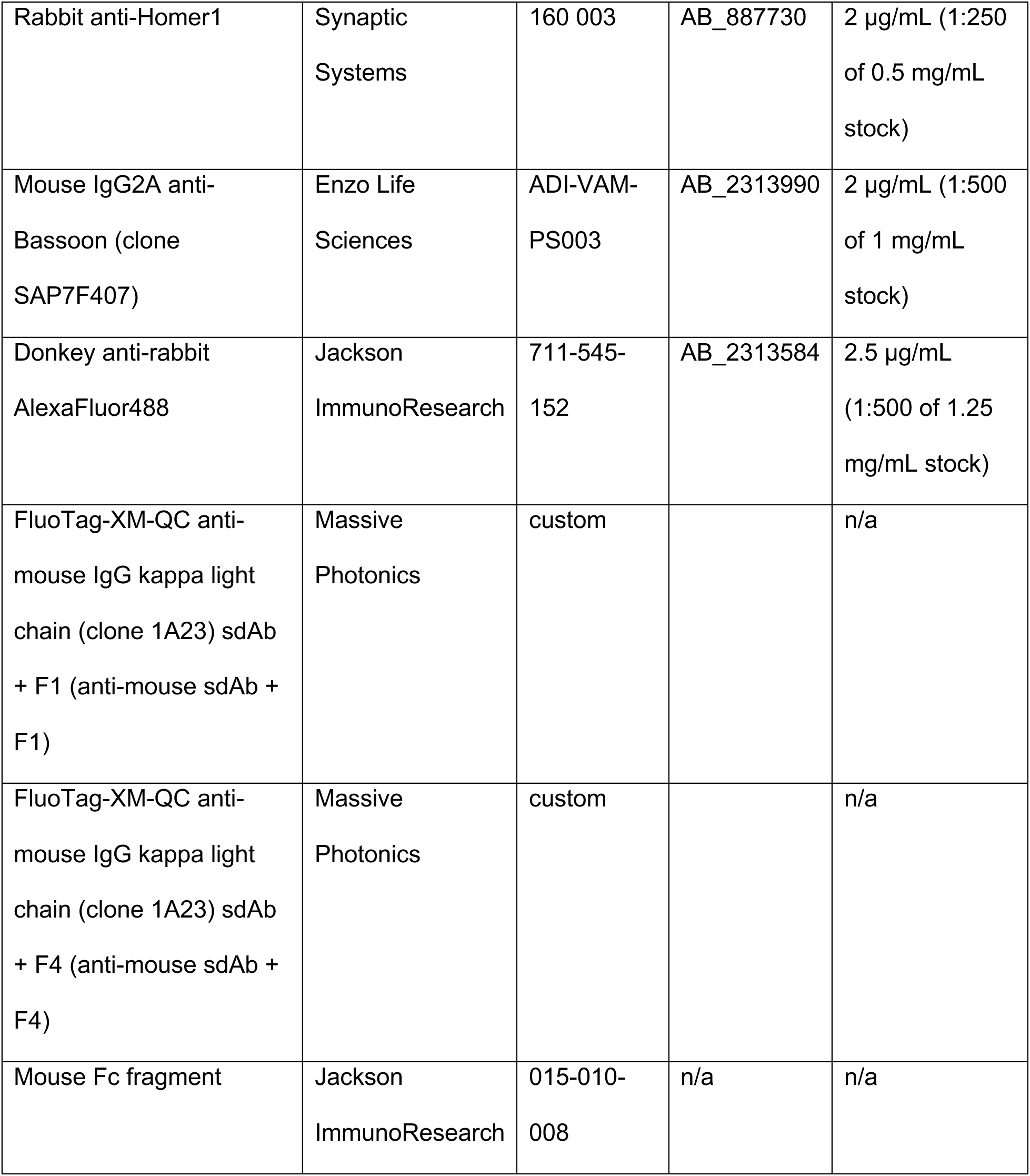

Common chemical reagents were purchased from Sigma. Reagents for staining included 16% EM grade paraformaldehyde (Electron Microscopy Sciences 15710), Triton X-100 (Sigma X100), donkey serum (Sigma D9663), and BSA (Sigma A7906). Antibodies used were mouse IgG2a anti-PSD-95 (Antibodies Inc 75-028, RRID: AB_2877189), mouse IgG1 anti-SAP102 (BioLegend 832001, RRID: AB_2564952), rabbit anti-Homer1 (Synaptic Systems 160 003, RRID: AB887730), donkey anti-rabbit AlexaFluor488 (Jackson ImmunoResearch 711-545-152, RRID: AB2313584), and mouse Fc fragment (Jackson ImmunoResearch 015-010-008). FluoTag-XM-QC anti-mouse IgG kappa light chain (clone 1A23) conjugated to either docking strand F1 or to docking strand F4 (aka anti-mouse sdAb + F1/F4), as well as F1-Cy3B and F4- Atto643 imager strands, were custom-ordered from Massive Photonics. The orthogonal F1 and F4 docking strands were derived from Strauss & Jungmann, 2020.

### Antibody preincubation

Multiplexed staining for DNA-PAINT was achieved by preincubating primary antibodies with secondary sdAbs conjugated to DNA docking strands, as in Sograte-Idrissi et al., 2020. Mouse anti-PSD-95 and mouse anti-SAP102 were separately mixed with anti-mouse sdAb + F4 or F1, respectively, at a 2.5-fold molar excess of sdAb to IgG for 20 minutes at room temperature, which will saturate the binding sites of the monovalent sdAbs to the two copies of light chain per IgG. Preincubation mixes were then combined and diluted to their final staining concentrations as described below.

### Immunostaining

Cultured neurons were fixed with 2% PFA/4% sucrose in PBS for 8 minutes at room temperature (RT), then washed 3 x 5 minutes with PBS + 100 mM Glycine (PBS/Gly), permeabilized with 0.3% Triton X-100 in PBS/Gly for 20 minutes at RT and blocked with 10% donkey serum, 3% BSA, and 0.2% Triton X-100 in PBS/Gly for 60 minutes at RT. The cells were stained overnight at 4˚C with antibodies against mouse anti-PSD-95 (6.25 µg/mL final concentration) and mouse anti-SAP102 (10 µg/mL final concentration) that were pre-incubated with anti-mouse sdAb F4 or F1, respectively, for DNA-PAINT, as well as with rabbit anti-Homer1 (2 µg/mL final concentration) to identify synapses. Primary antibodies were diluted in 5% donkey serum, 1.5% BSA, 0.1% Triton X-100 in PBS/Gly. The next day, the cells were washed with PBS/Gly, incubated 1h RT with donkey anti-rabbit AlexaFluor488 (2.5 µg/mL final concentration) diluted in PBS/Gly, washed, post-fixed with 4% PFA/4% sucrose in PBS for 15 minutes, and washed a final 3 times before storage at 4˚C until imaging. For experiments with bassoon staining, cells were fixed in 2% PFA in 10mM MES (pH6.8), 138mM KCl, 3mM MgCl_2_, 2mM EGTA, 320mM sucrose) for 15 min, then stained with the same protocol, with either the PSD-95 or SAP102 antibody replaced with mouse anti-Bassoon (2 µg/mL final concentration) preincubated with the appropriate sdAb.

### Microscope setup for DNA-PAINT

DNA-PAINT images were acquired on an Olympus IX81 inverted microscope with an Olympus 100x/1.49 NA TIRF oil immersion objective. An arc lamp provided epifluorescence for identifying Homer1-stained regions of interest by eye, and excitation lasers from an Andor ALC (405/488/561) and a Toptica iBeam Smart (640) were reflected to the sample through a 405/488/561/638 quadband polychroic (Chroma) in Highly Inclined and Laminated Optical (HILO) illumination to maximally illuminate the sample plane while limiting background. Emission was passed through an adaptive optics system (MicAO, Imagine Optics) to correct aberrations in the point-spread function, then split by a DV2 image splitter (Photometrics) containing a T640lpxr dichroic as well as ET655lp single band (far-red) and 59004m dual-band (red and green) emission filters to allow for identification of Homer1-stained synapses (AlexaFluor488) and simultaneous collection of Cy3B and Atto643-labeled imagers without changing the optical configuration. Emission was collected with an iXon+ 897 EM-CCD camera (Andor). Z stability was maintained by an Olympus ZDC2 feedback positioning system, and the microscope was contained inside an insulated box with temperature control to minimize drift. All components were controlled by iQ3 software (Andor), except the Toptica laser (TOPAS iBeam Smart GUI) and the MicAO (Imagine Optic software).

### Single-molecule imaging

Regions of interest were identified by Homer1 staining. PSD-95 and SAP102 localizations were obtained by simultaneous imaging of F1-Cy3B and F4-Atto643 imager strands diluted to 1 nM in imaging buffer (1x PBS + 500 mM NaCl + oxygen scavengers (PCA/PCD/Trolox) (Schnitzbauer et al. 2017), with laser power densities out of the objective of ∼3.3 and ∼2.2 kW/cm^2^ for the 640 and 561 lasers, respectively, for 50,000 frames with 50 ms exposure. Separately, 8-10 fields of TetraSpeck beads (100 nm; Invitrogen T7279) immobilized on coverslips coated with poly-L-lysine were imaged for 100 frames with 50 ms exposure and low laser power for determining dual-view correction.

### Single-molecule image processing

Single-molecule movies were converted to .raw using the imagej-raw-yaml-export plugin (https://github.com/jungmannlab/imagej-raw-yaml-export) in FIJI (Schindelin et al. 2012), then localized and processed with a custom MATLAB script and Picasso command line calls (Schnitzbauer et al. 2017). First, spots for PSD-95 and SAP102 were localized separately using Picasso *localize* with minimum net gradient of 25000 for PSD-95 and 35000 for SAP102. Next, the localizations were drift corrected using redundant cross-correlation with Picasso *undrift* at 3000 frames per segment and then recombined to one image. Reference TetraSpeck bead images were localized, and localization clusters were paired between the two sides of the image and used to generate a polynomial transformation using MATLAB’s *fitgeotrans* function with a second order polynomial. This transform was used to correct chromatic aberrations between the synapse images using MATLAB *transformPointsInverse*, and any remaining offset between the two channels was corrected by calculating a linear shift to maximize the cross-correlation between the images. Localizations were then removed if they had: photon count less than the mode photon count, standard deviation of the fit greater than 1.5 pixels in x and y, or localization error greater than 20 nm in x and y. Background non-clustered localizations were removed using Picasso *dbscan* with a radius of 48 nm and minpts = 10, and small clusters (<75 locs) were eliminated. Finally, clusters that are likely to be non-specific binding are removed based on the cluster’s localization kinetics, as non-specific binding tends to occur for very few frames while real binding sites have localizations that occur throughout the entire acquisition. Clusters with localizations less than the mean frames multiplied by two times the standard deviation and outside of the standard deviation range of 500 to 2000 are removed. The remaining clusters are putative synapses.

### Synapse selection

Synapses from each protein undergo separate synapse selection and then are paired post-selection and filtering. Synapses are selected from the list of putative synapses in a series of filtering steps. First, putative synapses are manually screened rejected if they show artificially sparse or dense localization density, or if multiple synaptic puncta are detected as a single cluster. Next, an additional area cutoff is applied. This cutoff was determined from a separate experiment where the presynaptic marker bassoon was labeled along with SAP102 and PSD-95 (Fig. 1-1A, D). From that data, we determined the area of SAP102 or PSD-95 clusters that did and did not co-localize with bassoon, and used these distributions to determine an area cutoff for likely true synaptic MAGUK clusters (Fig. 1-B1, E). When clusters smaller than the cutoffs were rejected (SAP102 cutoff: 0.0135 mm^2^; PSD-95 cutoff: 0.0195 mm^2^), 85.2% of SAP102 puncta and 87.7% of PSD-95 puncta contained bassoon versus 80.3% and 82.2% when there is no cutoff (Fig. 1-1C, F). The remaining PSD-95 and SAP102 clusters were considered true synaptic puncta.

**Figure 1:**
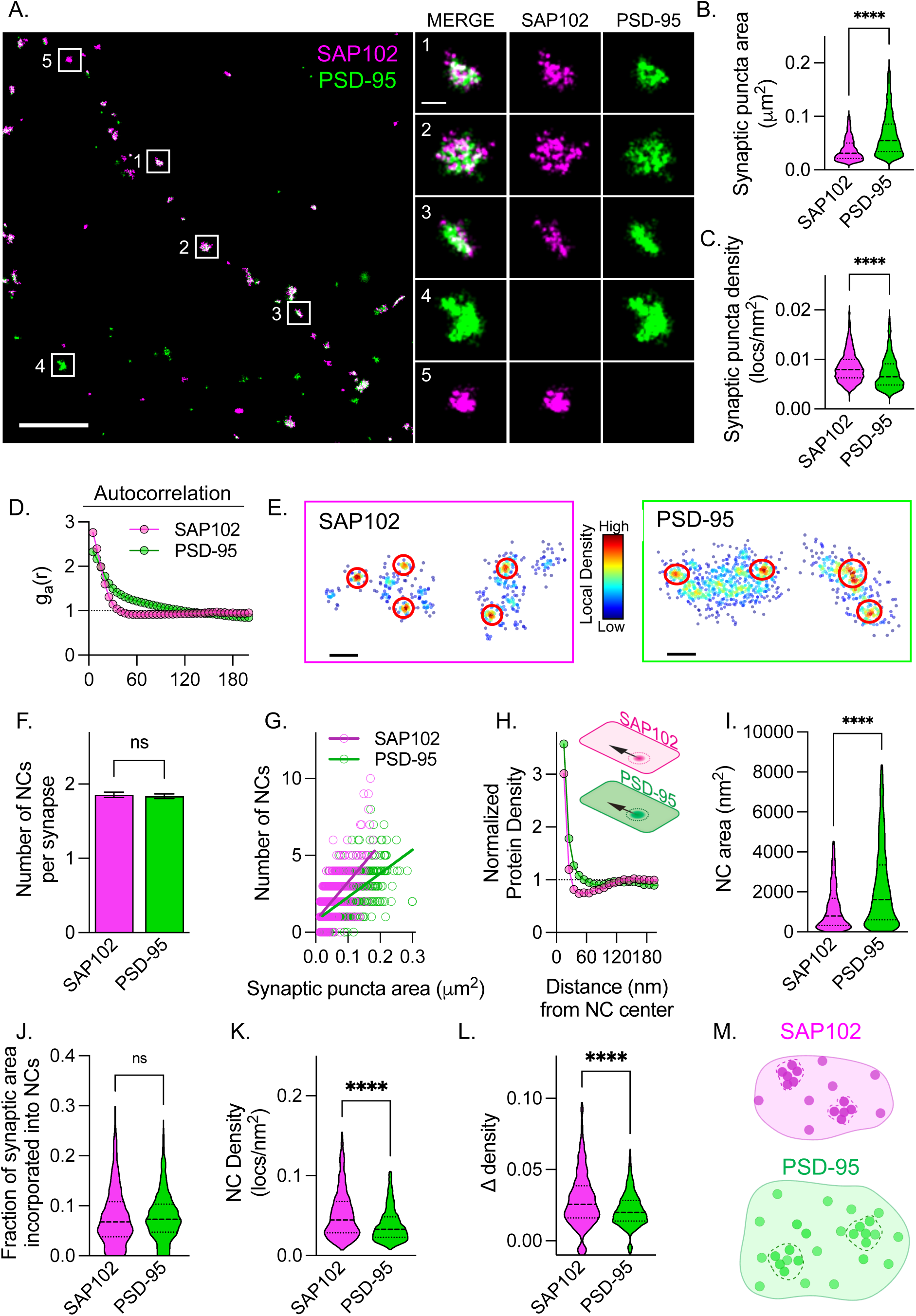
**SAP102 is organized into high-density subsynaptic nanoclusters with distinct properties.** Outliers removed to aid visualization for violin plots, full data set is used for statistics A. (Left) Example stretch of DNA-PAINT super-resolved dendrite with both SAP102 (magenta) and PSD-95 (green). Scale bar: 2.5 mm (Right) Magnified views of selected synaptic puncta indicated by white boxes. Synapses 1-3 contain both SAP102 (magenta) and PSD-95 (green) while synapse 4 contains only PSD-95 and synapse 5 contains only SAP102. Scale bar: 200 nm. B. Synaptic puncta area for SAP102 and PSD-95. (SAP102: 0.044 ± 0.001 mm^2^, n=1057 synapses; PSD-95: 0.071 ± 0.001 mm^2^, n=1293 synapses; p<0.0001, unpaired *t*-test) C. Overall synaptic puncta localization density for SAP102 and PSD-95. (SAP102: 0.009 ± 0.0001 locs/nm^2^, n=1057 synapses; PSD-95: 0.007 ± 0.0001 locs/nm^2^, n=1293 synapses; p<0.0001, unpaired *t*-test) D. Autocorrelation of SAP102 and PSD-95 distributions. E. Example localization maps for SAP102 (left) and PSD-95 (right). Colormap indicates local density around each localization. Red circles indicate high density areas that were identified as NCs based on the NC detection algorithm used. F. Number of NCs per synapse for SAP102 and PSD-95. (SAP102: 1.856 ± 0.0366, n=1057 synapses; PSD-95: 1.837 ± 0.0309, n=1293 synapses; p=0.684, unpaired *t-*test) G. Relationship between number of NCs and synaptic puncta area for SAP102 and PSD-95 (SAP102: slope=24.82, PSD-95: slope=15.39, p<0.0001). H. Autoenrichment analysis for both SAP102 and PSD-95. I. NC area based on the NC detection algorithm for SAP102 NCs and PSD-95 NCs. (SAP102: 1953 ± 62.95 nm^2^, n=1962 NCs; PSD-95: 3197 ± 79.54 nm^2^, n=2375 NCs; p<0.0001, unpaired *t*-test) J. Fraction of the synaptic puncta occupied by NCs for SAP102 and PSD-95. (SAP102: 0.081 ± 0.002, n=1057 synapses; PSD-95: 0.08 ± 0.001, n=1293 synapses; p<0.0001, unpaired t test) K. Internal NC localization density for SAP102 and PSD-95. (SAP102: 0.058 ± 0.001 locs/nm^2^, n=1962 NCs; PSD-95: 0.042 ± 0.0006 locs/nm^2^, n=2375 NCs; p<0.0001, unpaired *t-*test) L. The difference between the overall synaptic density and NC internal density (Ddensity) for SAP102 and PSD-95. (SAP102: 0.033 ± 0.001 locs/nm^2^, n=1057 synapses; PSD-95: 0.022 ± 0.0004 locs/nm^2^, n=1293 synapses; p<0.0001, unpaired *t-*test) M. Schematic representing the key features in the subsynaptic organization of SAP102 (magenta) and PSD-95 (green). Circles represent molecules of each protein, the overall synaptic puncta area is defined by the solid line, and NCs boundaries are indicated by dashed lines.

We next determined for each synaptic puncta whether it was colocalized with the other protein. Using custom MATLAB code, for every PSD-95 synaptic puncta, we asked whether there was a SAP102 synaptic puncta within a 500 nm box of the synapse edge. If a SAP102 puncta is present within the box and there is at least 30% area overlap between synaptic puncta, then they were categorized as paired. We found that decreasing this area overlap percentage cutoff further did not impact which puncta were paired (data not shown). Otherwise, the synaptic puncta are categorized as either SAP102 or PSD-95 alone.

### Synaptic analyses

Analyses of synaptic measures of DNA-PAINT data were done with custom MATLAB code largely as described (Chen et al., 2020), adapted for 2D localizations. Synaptic puncta area is determined by identifying the synapse border using the MATLAB *alphaShape* function. Synaptic density is calculated from the number of protein localizations divided by the area determined from the alpha shape. Autocorrelation analysis was performed as described previously (Tang et al. 2016; Ramsey et al. 2021).

### Nanocluster analysis

Previously our lab has detected NCs using custom MATLAB code (Chen et al., 2020; Tang et al., 2016). Due to the unique distribution of SAP102 we tested whether this approach was suitable for detection of nanoclusters of both SAP102 and PSD95 imaged by DNA PAINT. We compared a variety of parameters using our previously published approach and in addition two alternative cluster definition algorithms (DBSCAN (Dharmasri, 2023) and ToMATo (Pike et al., 2020)), and observed that all key relationships reported here were robust to most visually reasonable automated cluster definition strategies. We found that adjusting two key parameters of our previously published approach improved the detection of the NCs for SAP102 while maintaining detection of PSD-95 NCs and therefore was the best option. First parameter was the radius variable which defines the search area for neighboring localizations and second was the *cutoff* distance, which determines how far away a localization can be from an NC localization and still be considered part of that NC. We found at a radius variable of 2.6 and cutoff of 90 nm reliably detected NCs and defined their borders for both proteins. NC cluster-based measurements were made using custom MATLAB code essentially as described, with modifications for 2d (Tang et al., 2016). Autoenrichments were calculated as the density of localizations around each NC center normalized to a random distribution of localizations, as a function of distance from the NC center. NC area was determined from a convex hull of the NC localizations. NC density was calculated by dividing the number of localizations in each NC by its area. To measure the fraction of each synapse’s area occupied by NCs, the sum of the areas of all NCs in a synapse was divided by the synapse area. Ddensity was calculated by first finding the mean NC density for all NCs in a given synapse, then subtracting that from the overall localization density of the synaptic puncta. NC cross-enrichments were measured as for autoenrichments, but from the center of one protein’s NC to the density of the other protein. To calculate the area-scaled cross-enrichment, the radius of each NC was determined based on its area and assuming the NC is circular, and a scaling factor of 1.5 is applied to radius (radius*1.5) as a buffer to account for lack of circularity for all NCs and any possible inaccuracies in determining NC boundaries. Normalized protein density is then averaged for the cross-enrichment bins spanning the scaled radius (rounded up), to get an area-scaled cross-enrichment per NC.

### Statistics

All plots were generated and statistics performed in Prism (Graphpad). For violin plots, outliers were removed using ROUT to aid visualization, but statistics were performed on the full datasets without outlier removal. To make comparisons when only two groups were present, an unpaired t-test was used. For comparison of SAP102 and PSD-95 in alone vs paired synapses as well as across development, a two-way ANOVA with Tukey’s multiple comparisons post-hoc test was used. All experiments were repeated in 3 independent cultures. In text and figure legends, statistics are reported as mean ± SEM.

## Results

### SAP102 forms subsynaptic nanoclusters with distinct properties

We utilized DNA-PAINT super-resolution imaging to co-visualize the subsynaptic nano-organization of SAP102 and PSD-95. DNA-PAINT is a single-molecule localization microscopy method that allows for nanometer-scale resolution imaging of multiple proteins simultaneously (Friedl et al., 2023; Schnitzbauer et al., 2017) while avoiding complications due to fluorophore photophysics such as in dSTORM. We fixed cultured rat hippocampal neurons at DIV13-15, a developmental timepoint with significant expression of both SAP102 and PSD-95 (Cizeron et al., 2020), and immunolabeled endogenous PSD-95 and SAP102 for DNA-PAINT. In order to select puncta that likely represent synaptic versus non-synaptic immunolabeling, we determined size cutoffs in a separate experiment that included presynaptic bassoon labeling. We chose area cutoffs such that for the selected puncta at least 85% were associated with a presynaptic marker and therefore are likely synaptic (Fig. 1-1A-F) Using this approach, we were able to reliably super-resolve SAP102 and PSD-95 at individual synapses (Fig. 1A; left). Consistent with previously published observations, we found that while many synapses contained both SAP102 and PSD- 95, a subset contained only one protein or the other (Cizeron et al., 2020) (Fig. 1A; right). Considering each protein’s synaptic puncta independently, we found that SAP102 synaptic puncta were smaller than PSD-95 (0.044 ± 0.001 mm^2^ vs 0.071 ± 0.001 mm^2^; Fig.1B) and denser (0.009 ± 0.0001 locs/nm^2^ vs 0.007 ± 0.0001 locs/nm^2^; Fig.1C) than those of PSD-95. Using autocorrelation analysis, an unbiased approach for assessing density patterns (Tang et al., 2016; Veatch et al., 2012), we observed that SAP102 and PSD-95 both had autocorrelation functions that started well above 1, consistent with a non-random subsynaptic distribution (Fig. 1D). This indicates that both proteins are arranged non-homogeneously within the synapse, as was described for PSD-95 previously (Fukata et al., 2015; MacGillavry et al., 2013; Nair et al., 2013). However, the SAP102 autocorrelation curve had some key differences compared to PSD-95, including a higher initial peak and steeper early slope, suggesting distinct subsynaptic organizational principles between the two MAGUKs. Based on the higher peak and steeper slope, we predicted that SAP102 molecules accumulated into smaller subsynaptic areas of higher density relative its synaptic density compared to PSD-95, while the lower valley following the peak suggests sparser SAP102 between its high-density areas compared to PSD-95.

To describe SAP102 accumulation into high-density regions and compare its nano-organization to that of PSD-95, we used an automated algorithm to detect subsynaptic nanoclusters (NCs) of each protein (Chen et al., 2020; Ramsey et al., 2021; Tang et al., 2016). We used this approach to measure nanocluster features for each protein, and to characterize SAP102 nanostructure in contrast to that of PSD-95. Each protein had a similar number of NCs per synaptic punctum (SAP102: 1.856 ± 0.036 vs PSD-95: 1.837 ± 0.031; Fig. 1F). However, given the smaller overall area of SAP102 puncta, we asked whether the relationship between NC number and puncta area was different between SAP102 and PSD-95. We found that for both proteins, the two features were correlated (SAP102: R^2^= 0.416, p<0.0001; PSD-95: R^2^=0.439, p<0.0001; Fig. 1G), yet the slopes of these relationships were significantly different from one another (p<0.0001; Fig. 1G). This suggests that how MAGUK NCs are established is different for SAP102 and PSD-95, and that while overall NC number scaling with puncta area may be a conserved feature of MAGUKs (Broadhead et al., 2016), there appears to be an additional underlying mechanism at individual synapses that modifies the number of scaffold protein NCs independent of each MAGUKs overall content.

We next measured the normalized density of each protein with respect to the center of its own NC (autoenrichment), which gives insight into the size of NCs and how they relate to the space around them (Fig. 1H). We found that like the autocorrelation, SAP102 and PSD-95 had high-magnitude normalized protein densities close to their NC centers. However, away from this center, SAP102 density declined to average levels (a normalized protein density of 1) within a shorter distance than did PSD-95, indicating that SAP102 NCs are smaller than PSD-95 NCs (SAP102: between 25 and 35 nm away from NC center; PSD-95: between 55 and 65 nm away from NC center; Fig. 1H). Additionally, unlike PSD-95, SAP102 normalized protein density declined from its peak to values less than 1, indicating SAP102 is in fact de-enriched outside of its NCs (Fig. 1H). This was borne out in the analysis of the size and features of individual NCs following the definition of their borders by applying a local density cutoff for each localization (see methods). SAP102 NCs were smaller than those of PSD-95 (SAP102: 1953 ± 62.95 nm^2^; PSD- 95: 3197 ± 79.54 nm^2^, Fig. 1I) and the total area per synapse occupied by NCs was not different between proteins (SAP102: 0.081 ± 0.002; PSD-95: 0.08 ± 0.001; Fig. 1K). SAP102 NCs were also more internally dense (SAP102: 0.058 ± 0.001 locs/nm^2^ vs PSD-95: 0.042 ± 0.0006 locs/nm^2^; Fig. 1J). To assess the propensity of each protein to accumulate within NCs at each synapse, we calculated the difference between the average density within its NCs and its density overall within the synaptic punctum (Δdensity). Unlike for area, the relationship between NC density and overall synaptic density was different between SAP102 and PSD-95. SAP102 puncta had a larger Δdensity (0.033 ± 0.001) than PSD-95 puncta (0.023 ± 0.0004; Fig. 1L), consistent with less enrichment of SAP102 outside of NCs than is seen for PSD-95.

Overall, these data reveal that SAP102, like PSD-95, has a heterogeneous subsynaptic distribution typified by organization into several high-density NCs within the bounds of the PSD overall. However, SAP102 tends to form smaller NCs with less protein between them than does PSD-95 (Fig. 1M).

### Nano-organization at synapses lacking either SAP102 or PSD-95 is distinct from synapses with both proteins

It has not been clear how the organization of MAGUK proteins are influenced by the presence of multiple family members. To address this, we noted that while many synapses have both SAP102 and PSD-95, a subset contained only one of the proteins. Thus to test whether SAP102 or PSD-95 nanoscale organization was influenced by the presence of the other protein, we divided the synapses into 3 categories: SAP102 alone, PSD-95 alone, and paired synapses which contained both proteins. For both proteins, the area bounded by the protein was smaller at synapses that contained only one protein versus both (Table 2-1; Fig. 2A) suggesting an interaction. The autocorrelation curves for SAP102 and PSD-95 displayed similar overall characteristics whether the proteins were paired or alone (Fig. 2C) indicating that in both synapse types the proteins form subsynaptic NCs. However, the curves for each protein began to plateau at a lower autocorrelation value when alone than when paired. SAP102 and PSD-95 autoenrichment analysis mirrored the autocorrelation curves (Fig. 2D). The concurrence between these two measures suggests that in the absence of the other protein, SAP102 and PSD-95 tend to be more enriched within nanoclusters, despite forming smaller synaptic puncta.

**Figure 2:**
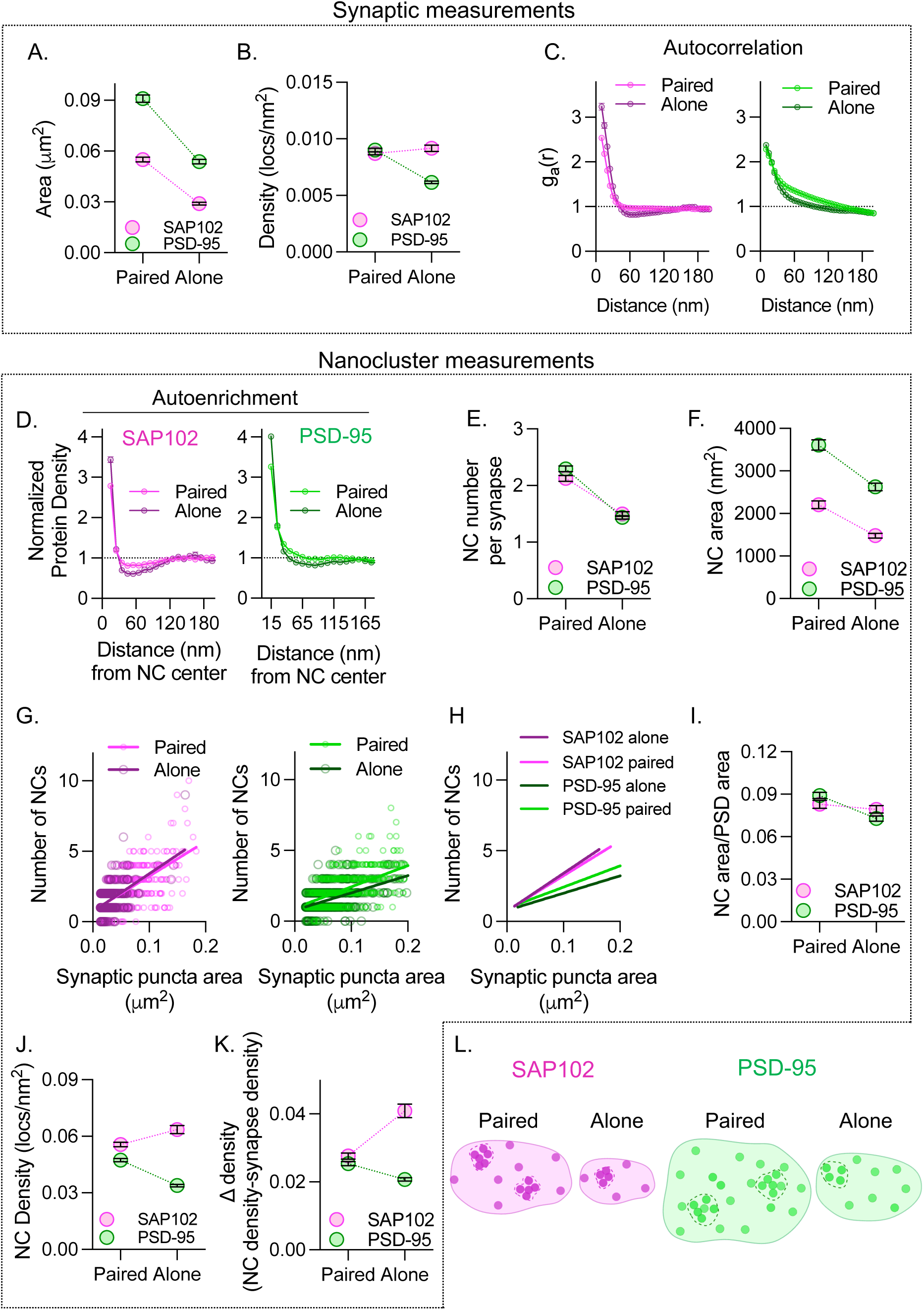
**SAP102 and PSD-95 nanostructure is impacted by the presence of the other protein** A. Synaptic puncta area for SAP102 (magenta) and PSD-95 (green) at synapses with both proteins present (paired) or just one of the proteins (alone) (See Table 2-1 for statistics). B. Synaptic puncta density for SAP102 and PSD-95 at paired and alone synapses (See Table 2-1 for statistics). C. Left: autocorrelation analysis for SAP102 at paired (light magenta) and alone (dark magenta) synapses. Right: PSD-95 autocorrelation analysis at paired (light green) and alone (dark green) synapses. D. Autoenrichment analysis for SAP102 at paired and alone synapses (left) and PSD-95 at paired and alone synapses (right). E. Number of NCs per synapse for SAP102 and PSD-95 at paired and alone synapses (See Table 2-1 for statistics). F. NC area for SAP102 and PSD-95 at paired and alone synapses (See Table 2-1 for statistics). G. Relationship between number of NCs and synaptic puncta area for SAP102 (left; SAP102 paired: slope=24.62, SAP102 alone: slope=26.82, p=0.417) and PSD-95 (right; PSD-95 paired: slope=15.00, PSD-95 alone: slope=12.24, p=0.0106) at both paired and alone synapses. H. Linear regression lines describing the relationship between NC number and synaptic puncta area for SAP102 and PSD-95, reproduced from 2G for clarity. I. Area of the synapse incorporated into NCs for SAP102 and PSD-95 at paired and alone synapses (See Table 2-1 for statistics). J. NC internal density for SAP102 and PSD-95 at paired and alone synapses (See Table 2-1 for statistics). K. Ddensity for SAP102 and PSD-95 at paired and alone synapses (See Table 2-1 for statistics). L. Schematic representing the differences in subsynaptic organization between SAP102 (magenta) alone vs paired and PSD-95 (green) alone vs paired. Circles represent molecules of each protein, the overall synaptic puncta area is defined by the solid line, and NC boundaries are indicated by dashed lines.

SAP102 and PSD-95 each had fewer, smaller NCs at synapses where they are alone versus paired (Table 2-1; Fig. 2E-F). Additionally, NC number and synaptic puncta area were correlated for both proteins whether alone (SAP102 alone: R^2^=0.282, p<0.0001; PSD-95 alone: R^2^=0.294, p<0.0001; Fig. 2G) or paired (SAP102 paired: R^2^=0.404, p<0.0001; PSD-95 paired: R^2^=0.403, p<0.0001; Fig. 2G). However, while the relationship between NC number and area was consistent for both categories of SAP102 synapses (paired slope= 24.62, alone slope= 26.82, p=0.417; Fig. 2G), there was a difference between slopes for PSD-95 paired vs alone (paired slope= 15.00, alone slope= 12.24, p=0.0106; Fig. 2G). Thus the higher number of PSD-95 NCs at paired synapses was not due simply to the increase in overall synaptic puncta area, but instead the presence of SAP102 was associated with a higher propensity of PSD-95 to form NCs (Fig. 2H). Interestingly, the proportion of total protein area per puncta occupied by NCs was maintained for SAP102 whether alone or paired (Table 2-1), whereas, for PSD-95 the total NC area normalized to its puncta area was smaller for PSD-95 alone than paired (Table 2-1; Fig. 2I). When we compared NC properties between proteins, we found that the differences between PSD-95 and SAP102 nano-organization persisted regardless of whether they were paired or alone, with SAP102 tending to have smaller NCs than PSD-95, and no difference in overall NC number or proportion of area occupied by NCs between the two proteins (Table 2-1; Fig. 2E-H).

When we compared density measures between synapses with both proteins and those with just one, we found that there were substantial differences between SAP102 and PSD-95. Compared to paired synapses, SAP102 density when alone was not different whereas PSD-95 density was lower when alone (Table 2-1; Fig. 2B). This may reflect that SAP102 recruitment to synapses is independent of PSD-95, whereas the two work synergistically to recruit additional PSD-95. Additionally, at synapses with both proteins, SAP102 and PSD-95 synaptic densities were not different from each other, while at alone synapses, SAP102 density was higher than PSD-95 (Table 2-1; Fig. 2B). Furthermore, SAP102 NC density was higher in SAP102 alone synapses than in paired synapses, whereas PSD-95 NC density was lower (Table 2-1; Fig. 2J). Finally, at paired synapses, Δdensity was the same for both proteins, while at synapses with each protein alone, SAP102 had a much higher Δdensity than PSD-95 (Table 2-1; Fig. 2K). These results indicate that in the absence of PSD-95, SAP102 has a higher tendency to accumulate within NCs than PSD-95 does alone, whereas when both are present their density within NCs is more matched, perhaps due to space constraints or subsynaptic densities of interacting proteins.

Since SAP102 and PSD-95 compensate for at least a subset of one another’s functions following knockout (Cuthbert et al., 2007), we had predicted that subsynaptic organizational properties would converge to a common form at synapses containing only one scaffold, whereas the differences between their nanoscale organization would be enhanced when both are present. However, these data reveal that rather than converge into similar structures when alone, SAP102 and PSD-95 nanostructural differences persisted, and in the case of NC density, their differences were actually enhanced when alone (Fig. 2L).

### SAP102 and PSD-95 occupy both overlapping and distinct subsynaptic nanodomains

We next determined how SAP102 and PSD-95 are arranged with respect to each other within individual synapses when both are present. We sought to distinguish between two general categories of potential spatial organizations. On one extreme, due to their high structural homology and shared binding partners, SAP102 and PSD-95 might overlap extensively within the synapse and form nanodomains with each present. On the other extreme, despite their similarities, their structural and interaction differences may be sufficient to promote their organization into separate domains, perhaps even tiling the synapse with unique functional subdomains.

Because we observed that the area of the synapse bounding SAP102 tended to be smaller than that encompassing PSD-95 even at synapses with both proteins, we first compared the contribution of SAP102 and PSD-95 to the overall synaptic puncta area at individual synapses. Total synaptic puncta area was determined by combining the areas bounded by SAP102 and PSD-95 (Fig. 3A: blue line). Consistent with SAP102 forming smaller overall synaptic puncta, we found that SAP102 occupied 52.8 ± 0.7% of the total synaptic area, while PSD-95 occupied 83.3 ± 0.6% (Fig. 3B). Of the total area, 41.3 ± 0.5% was occupied by both proteins (Fig. 3C), and thus the remaining portion of the synapse contained only one protein or the other. Of the total SAP102 area at the synapse, 79.5 ± 0.7% tended to overlap with PSD-95, while for PSD-95, only 49.9 ± 0.7% overlapped with SAP102 (Fig. 3C). Overall, these results show that when both are present, SAP102 covers less of the synapse and tends mostly to overlap with PSD-95, and that more of the PSD area contains PSD-95 alone than SAP102 alone.

**Figure 3:**
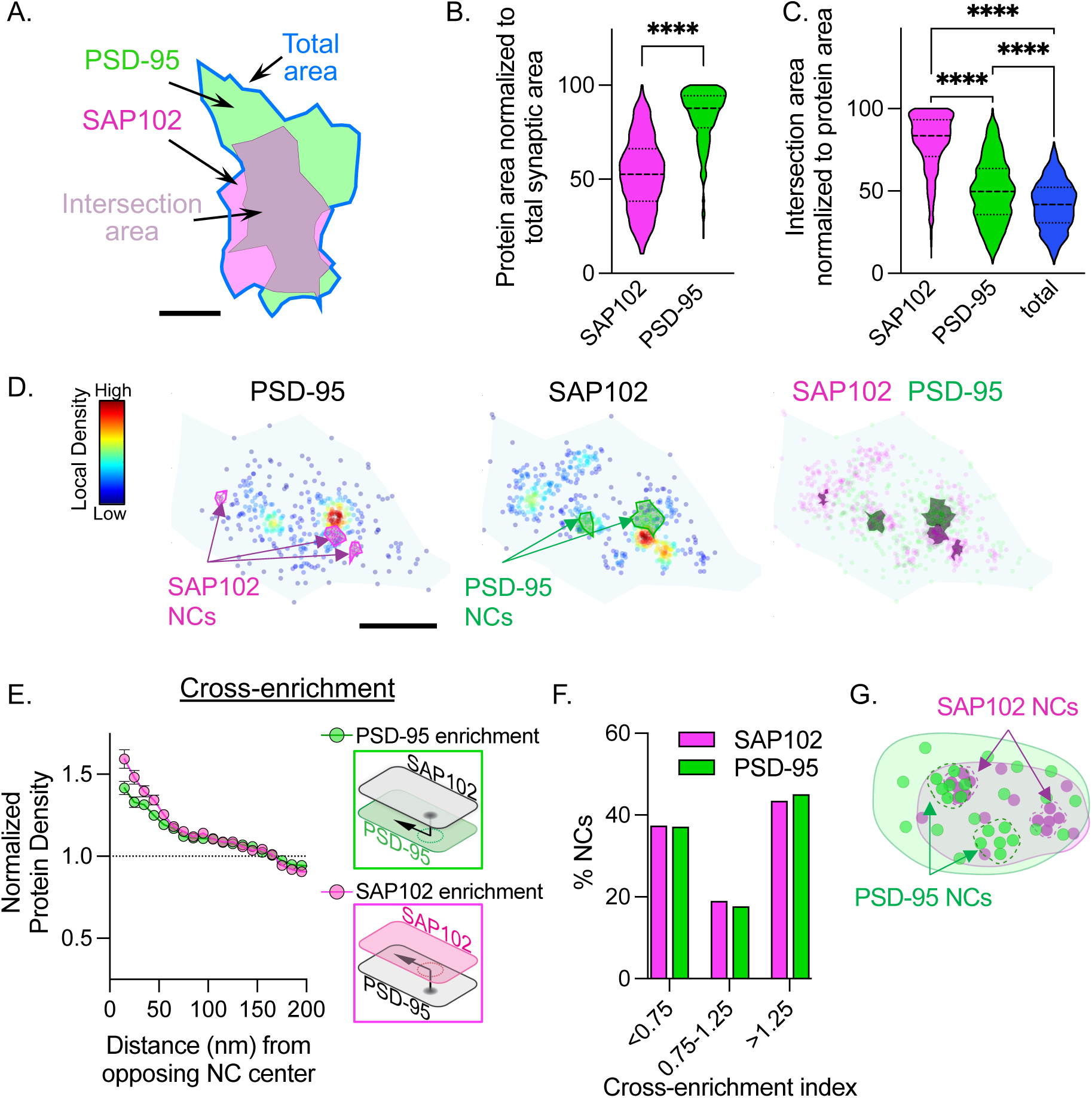
**SAP102 and PSD-95 occupy both distinct and overlapping subsynaptic nanodomains.** A. Example synapse containing both PSD-95 (green) and SAP102 (magenta). Gray area is the intersection area where the proteins’ synaptic puncta overlap. Blue line indicates the total synapse area, defined by overlaying and combining the SAP102 and PSD-95 synaptic puncta. Scale bar is 100 nm. B. Percentage of the total combined synaptic area occupied by SAP102 and PSD-95. Outliers removed to aid visualization for violin plots, full data set is used for statistics. (SAP102: 52.84 ± 0.783 % of total area, n=602 synapses; PSD-95: 83.33 ± 0.616 % of total area, n=602 synapses; p<0.0001, unpaired *t-*test) C. Percent of SAP102, PSD-95, and total synaptic areas that contain both SAP102 and PSD- 95 (intersection area). (SAP102: 79.53 ± 0.716 % overlap, n=602 synapses; PSD-95: 49.99 ± 0.785 % overlap, n=602 synapses; total: 41.36 ± 0.593 % overlap, n=602 synapses; p<0.0001, unpaired *t-*test) D. Example synapses with heat map indicating local density for each protein’s localizations (PSD-95: left and SAP102: middle) each with detected NCs for the other protein indicated (left: SAP102 NC in magenta; middle: PSD-95 NCs in green). Right: SAP102 (magenta) and PSD-95 (green) localizations with NCs for each protein overlaid. E. SAP102 and PSD-95 cross-enrichment with respect to NC centers of the other protein. Right: schematic illustrating the cross-enrichment analysis. F. Distribution of cross-enrichment indices scaled to NC area for each protein’s NCs. G. Schematic representing the distribution of SAP102 (magenta) and PSD-95 (green) within individual synapses. Circles represent molecules of each protein; the overall synaptic puncta area is defined by the solid line, and NCs boundaries are indicated by dashed lines.

To explore the relationship between the two proteins with regard to their NCs, we used a cross-enrichment analysis (Tang et al., 2016) to measure each protein’s density as a function of distance from the opposing protein’s NC center (Fig. 3D). SAP102 had a high relative density near PSD-95 NCs and vice versa (Fig. 3E), suggesting a substantial colocalization of the high-density domains of each protein. However, visual inspection suggested that while many SAP102 and PSD-95 NCs did colocalize with high-density regions of the other protein, others did not (Fig. 3D). To assess the proportion of NCs that were enriched with the other protein, for each NC we measured the opposing protein’s enrichment within an area proportional to the size of the NC (a value we termed the cross-enrichment index). This revealed that while the largest proportion of NCs of both proteins were cross-enriched for the other (cross-enrichment index >1.25; 43.5% of SAP102 NCs with PSD-95 and 45.1% of PSD-95 NCs with SAP102), nearly as many NCs of each had cross-enrichment indices below 0.75 (37.4% and 37.2% respectively), and this did not co-organize with the other protein (Fig. 3F). Thus, while a significant proportion of NCs do co-organize between proteins to form MAGUK nanodomains, there is also a large proportion that do not overlap within the same synapse. This suggests that when both proteins are present, these scaffolds establish several types of distinct postsynaptic nanodomains even within single synapses: those enriched for both, enriched only for one of these critical MAGUKs, and ones that fall somewhere in between where the protein NCs are near each other but not strongly co-localized. Given the known unique characteristics of SAP102 and PSD-95, these distinct MAGUK nanodomain types may contribute distinct synaptic functions.

### SAP102 and PSD-95 both undergo nanostructural changes across development, yet their relationship to each other is stable

SAP102 and PSD-95 each have well-described developmental profiles, with SAP102 peaking early in development on postnatal day 1 then declining, while PSD-95 synaptic expression peaks later at 3 weeks of age (Cizeron et al., 2020). Because their overall synaptic abundance and functional roles change over time (Elias et al., 2008), we asked whether their spatial organization evolves as synapses mature. To address this, we imaged neurons at one week in vitro (DIV 6-8) and three weeks in vitro (DIV 20-22) alongside the two-week-old neurons (DIV 13-15) described above (Fig. 4A). Consistent with previously published work, we observed an increase in synapses that contained both proteins or PSD-95 alone across development, while the percentage of synapses that contained only SAP102 decreased (Fig. 4B). Across development, the area of puncta containing SAP102 alone was relatively stable, while at paired synapses, the area bounded by SAP102 slightly increased (Table 4-1; Fig. 4C, left). This contrasts with PSD-95, for which the synaptic puncta area was larger in more mature synapses regardless of whether SAP102 was present (Table 4-1; Fig. 4C, right). Considering the area relationship between proteins at paired synapses, the fraction of the overall synapse occupied by SAP102 was higher at 3 weeks than at 1 and 2 weeks (Table 4-1; Fig. 4D, left), despite still only making up on average 57.6% of the total synapse area at 3 weeks in vitro. Consistent with the changes seen in PSD-95 synaptic puncta area, the area occupied by PSD-95 peaked at week 2 (Table 4-1; Fig. 4D, right). The area over which PSD-95 overlapped with SAP102 also peaked at week 2 (Table 4-1; Fig. 4E, left), while the overlap of SAP102 with PSD-95 peaked at week 3 (Table 4-1; Fig. 4E, right). Thus while PSD-95 consistently occupies a larger overall percentage of the synapse across development, the changing expression ratio of PSD-95 and SAP102 results in a shifting balance of the PSD territory they occupy. This extends previous work documenting how this shifting ratio alters the proportion of synapses containing only PSD-95 or SAP102 (Cizeron et al., 2020).

**Figure 4:**
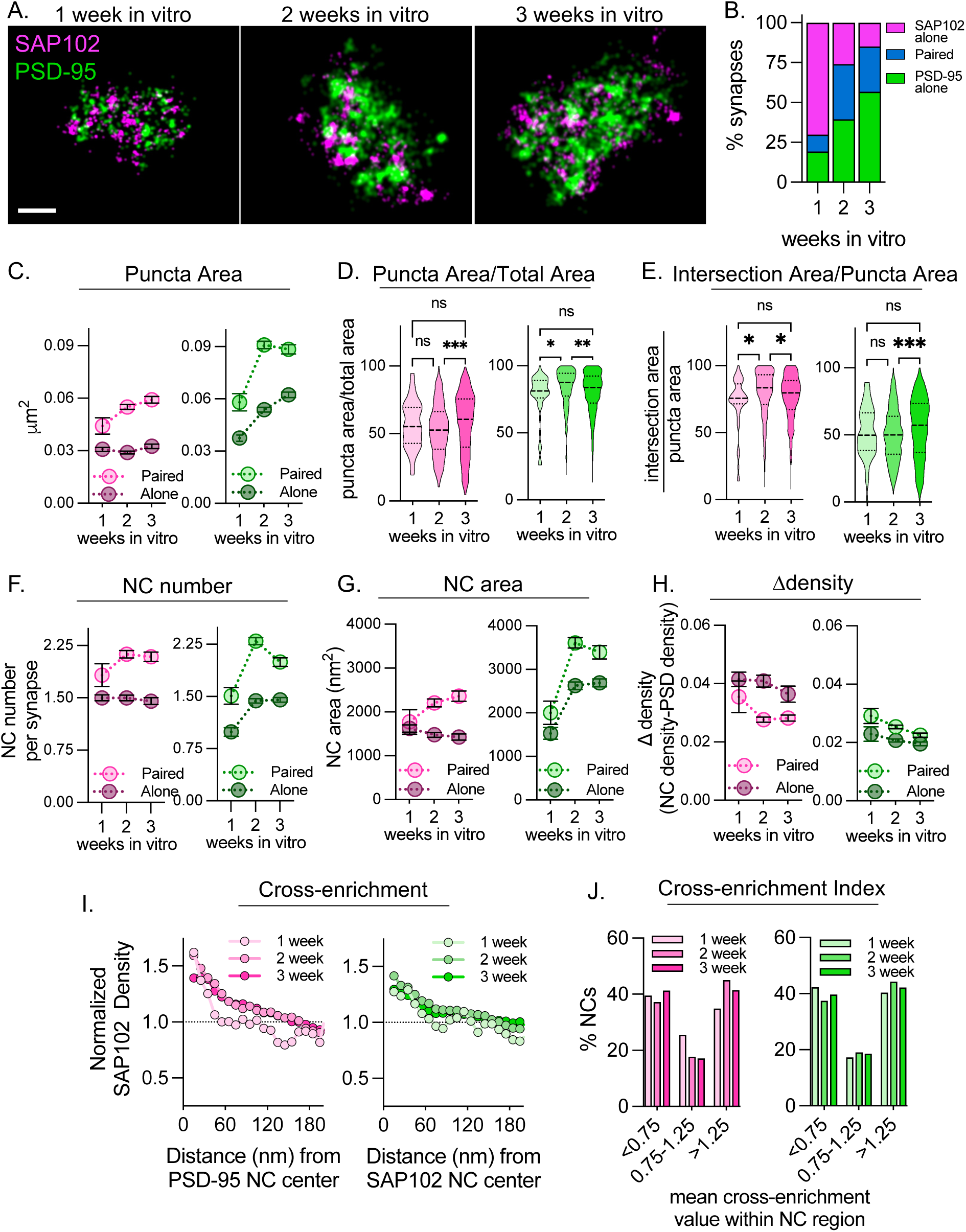
**SAP102 and PSD-95 nanostructure each change cross development, yet their nanoscale spatial relationship remains largely consistent.** *Throughout this figure-the 2 week in vitro dataset is the same dataset as in the previous figures. A. Example rendered synapses from 1, 2, and 3 weeks in vitro. Scalebar is 100 nm. B. Percentage of synapses with SAP102 alone (magenta), PSD-95 alone (green), or both proteins (blue) across development. C. Synaptic puncta area at both paired and alone synapses across development for SAP102 (left) and PSD-95 (right). D. Fraction of total synaptic puncta area occupied by SAP102 (left) or PSD-95 (right) across development. E. Fraction of each protein’s puncta area that overlaps with the other protein (intersection area) for SAP102 (left) or PSD-95 (right) across development. F. NC number at both paired and alone synapses across development for SAP102 (left) and PSD-95 (right). G. NC area at both paired and alone synapses across development for SAP102 (left) and PSD-95 (right). H. Ddensity at both paired and alone synapses across development for SAP102 (left) and PSD-95 (right). I. SAP102 (left) and PSD-95 (right) cross-enrichment with respect to NC centers of the other protein for both paired and alone synapses across development. J. Distribution of cross-enrichment indices scaled to NC area for each protein’s NCs for SAP102 (left) and PSD-95 (right) across development.

We next asked how each protein’s NC organization evolves over this period of rapid development. Overall, SAP102 tended to have a more stable nano-organization when alone rather than paired with PSD-95, showing little age-dependence of NC number, NC area or the density difference between the NC and the synapse overall (Table 4-1; Fig. 4F-H). In synapses with PSD-95 present, though, SAP102 NC number (Table 4-1; Fig. 4F, left) and NC area (Table 4-1; Fig. 4G, left) increased and Δdensity decreased (Table 4-1; Fig. 4H, left) across weeks. The number (Table 4-1; Fig. 4F, right) and area (Table 4-1; Fig. 4G, right) of PSD-95 NCs increased substantially after the first week, similar to what has been observed previously (Sun et al., 2022). Additionally, PSD-95 Δdensity did not change across development for alone or paired synapses (Table 4-1; Fig. 4H, right). This perhaps reflects that SAP102-alone synapses may represent an earlier phase in synaptic development, whereas when PSD-95 is also present, synapse maturation can proceed leading to changes in synaptic nanostructure.

Finally, we asked whether the nanoscale spatial relationship between SAP102 and PSD- 95 at paired synapses changes across development. An increase in subsynaptic segregation might suggest an increasing functional specialization. Interestingly, SAP102 and PSD-95 at 1 week did show a steeper initial slope in the cross-enrichment than the two later time points (Fig. 4I). This indicates that SAP102 at 1 week is enriched within a smaller radius from PSD-95 NC peaks and vice versa, perhaps due to the smaller size of PSD-95 NCs at this time point. Otherwise, there was overall little change in cross-enrichment between the 3 age groups for either protein. The overall proportion of NCs that were cross-enriched with had little variation across development for SAP102 or PSD-95 (Fig. 4J). Overall, the subsynaptic spatial relationship between SAP102 and PSD-95 was largely stable across these weeks of development despite changes in both overall synapse size and nanostructure and the shift in function and abundance of PSD-95 and SAP102 occur over the same timescale (Elias et al., 2008).

Taken together, the organization of SAP102 into unique subsynaptic NCs and the complex but developmentally persistent nanostructural relationship of SAP102 and PSD-95 suggest that subsynaptic spatial relationship may be a novel mechanism for functional differentiation of the MAGUK family.

## Discussion

The subsynaptic distribution of MAGUK proteins is of particular importance since they are central to the functional organization of synaptic signaling components. Using DNA-PAINT super-resolution imaging, we described the nanoscale organization of SAP102 within synapses and its relationship to PSD-95 nanostructure. We found that SAP102, like PSD-95, forms subsynaptic nanoclusters within individual synapses where protein is concentrated. However, SAP102 NCs have properties distinct from PSD-95 and tend to be smaller and denser with less protein outside of NCs. Most fundamentally, our data has revealed that SAP102 and PSD-95 are neither reliably segregated within synapses nor are they systematically co-localized within postsynaptic MAGUK nanodomains. This observation suggests that one role of different MAGUKs in the synapse may be to anchor synaptic proteins in specific postsynaptic nanodomains defined by the resident MAGUK. Their nanostructural diversity even when both are present, may reflect the formation of unique synaptic nanodomain subtypes that have distinct functional roles. Across the first weeks of cell development in vitro, while both proteins had some degree of nanostructural changes, the overall relationship between SAP102 and PSD-95 was preserved, suggesting that the spatial coordination between MAGUK protein nanostructure may be closely regulated.

Whether different MAGUK proteins help delineate different subdomains within synapses has not been explored previously. We found that in synapses that contain both PSD-95 and SAP102, ∼45% of the nanoclusters of either protein co-localize with the other. Thus, within individual synapses, there are nanodomains that contain SAP102 and PSD-95 clustered together at elevated concentrations, as well as nanoclusters of each protein relatively lacking the other which offers a platform for functional diversity within single synapses. Most obviously, this may underlie the ability of SAP102 and PSD-95 to serve both overlapping and distinct functions at synapses by positioning them in different proximities to their interactors. Both SAP102 and PSD- 95 can anchor AMPARs and NMDARs within the synapse (Elias et al., 2008; Lau & Zukin, 2007; Su et al., 2018; Zhu et al., 2016). However, there are differences in how each of these scaffolds interacts with receptors which may lead to differences in receptor composition at nanodomains that contain one versus both proteins. SAP102 prefers GluN2B-containing NMDARs (Sans et al., 2000), predicting accumulation of GluN2B-NMDARs in subsynaptic compartments enriched for SAP102, while GluN2A-containing receptors may co-organize with PSD-95. This might underlie the observation that NMDAR subtypes sometimes segregate into distinct subsynaptic areas (Kellermayer et al., 2018), though multiplexed labeling of NMDAR subtypes and MAGUKs will be required to test this. Interestingly, SAP102 mediates removal of NMDARs from the synapse in addition to their recruitment and stabilization (Chen et al., 2012). Because recruitment and elimination are opposing functions, it is possible that there is spatial segregation of these functions to discrete subsynaptic nanodomains.

The complexity of the SAP102 vs PSD-95 distribution may help establish a complex transsynaptic relationship between presynaptic release sites and postsynaptic receptor pools. While PSD-95 NCs align with presynaptic release sites and can anchor AMPA receptors to support their activation (Ramsey et al., 2021), more detailed 3-dimensional multiplexed imaging will be required to test whether these nanodomains contain both PSD-95 and SAP102, or whether SAP102-alone NCs are similarly aligned with or perhaps even positioned away from certain release sites. Interestingly, there is some evidence that within synapses there are distinct pools of receptors that respond to either spontaneous or evoked vesicle release events (Li et al., 2021; Reese & Kavalali, 2016; Sara et al., 2011; Wang et al., 2022). Segregation of MAGUK nanodomains suggests that these two scaffolds may separately facilitate alignment of receptors with distinct presynaptic release modes, for instance with SAP102 NCs enriching receptors away from evoked release sites while PSD-95 positions receptors near them to maintain distinct release mode-specific receptor pools.

While further exploration is required to determine the underlying origin of their similar yet distinct nanostructural organization, differences in SAP102 and PSD-95 post-translational modifications likely play a role. For one, the major isoforms of PSD-95 can be palmitoylated near the N-terminus, while SAP102 cannot (El-Husseini et al., 2000). This palmitoylation promotes targeting of PSD-95 to the synaptic membrane and is required for its synaptic accumulation and likely its organization into subsynaptic nanoclusters (Balderas et al., 2022; El-Husseini et al., 2002; Fukata et al., 2015; Schlüter et al., 2006; Zheng et al., 2010). SAP102, as it is non-palmitoylated, is found more often at sites farther from the synaptic membrane (Zheng et al., 2010, 2011), and its synaptic accumulation and stabilization is instead dependent on its SH3 and GK domains (Zheng et al., 2010). Interestingly, palmitoylation of PSD-95 is also tied to structural changes in its SH3/GK domain (Fukata et al., 2013), suggesting that membrane-proximal and distal interactions of these proteins may contribute differentially to their nano-organization. Other differential post-translation modifications between MAGUKs may also contribute to their subsynaptic clustering properties. Notably, synaptic accumulation of SAP102 is supported by phosphorylation at Ser632, whereas CaMKII-dependent phosphorylation of PSD-95 at Ser73 regulates stability of PSD-95 and other PSD residents including Shank2 (Steiner et al., 2008; Wei et al., 2018).

The distinctive distributions of PSD-95 and SAP102 may arise from their respective mobilities at synapses. Based on recovery after photobleaching, SAP102 is more mobile than PSD-95 (Zheng et al., 2010) as it recovers more quickly and has a smaller fraction of immobilized molecules in spines. Interestingly, the size of the mobile population of SAP102 but not PSD-95 is sensitive to pharmacological stabilization of the actin cytoskeleton (Zheng et al., 2010), suggesting that patterns of dynamic actin in contact with the PSD may specifically anchor SAP102 molecules in NCs. Finally, liquid-liquid phase separation (LLPS) of PSD-95 has been suggested to a play a role in subsynaptic organization (Feng et al., 2019; Liu et al., 2021; Zeng et al., 2016), though because direct observation of LLPS has relied on non-neuronal preparations, the extent of LLPS at synapses is ambiguous. Little is known about the propensity or characteristics of SAP102 LLPS, though recently, it was shown to be capable of forming condensates *in vitro* with the AMPAR binding partner stargazin (Zeng et al., 2019). Whether phase separation plays a role in determining subsynaptic spatial organization or whether SAP102 and PSD-95 form condensates together is not clear, though the higher mobility of SAP102 than PSD-95 (Zheng et al., 2010) is consistent with a higher partitioning into phase condensates as opposed to stably bound ensembles. We can speculate that one model for how MAGUK specific nanostructure is established is that palmitoylated PSD-95 creates a flexible but stabilized mesh with somewhat regular spacing (Chen 2008), within which nanoclusters of PSD-95 are created through interactions with abundant receptors and adhesion molecules. In contrast, SAP102 likely distributes subsynaptically primarily through multiple interactions of its PDZ domains and other domains in the cytosolic or “pallium” (Dosemeci et al., 2016) portion of the PSD, where it can more avidly undergo phase separation, interact with the actin cytoskeleton, and exchange readily with cytoplasmic pools in the spine and dendritic shaft.

Nanostructural properties of SAP102 and PSD-95 diverged at synapses with both proteins compared to those with either one alone. This was surprising, as one prediction is that the ability for compensation among MAGUKs (Won et al., 2017) would arise from similar nano-organization. The differences might reflect several possible mechanisms. One is that synapses lacking PSD- 95 or SAP102 may represent different stages of synaptic maturation. *In vivo* characterization has shown that the population of synapses without PSD-95 decreases across development while the population lacking SAP102 increases (Cizeron et al. 2020), consistent with our observations. Our results extend this observation by revealing that the balance of subsynaptic territory occupied by PSD-95 and SAP102 shifts with their changing expression levels, as opposed to simply shifting the proportion of single-MAGUK synapses. Therefore, it is possible that even at different ages, synapses containing only SAP102 or only PSD-95 are effectively at different maturation states and their unique nanostructure reflects their accordingly unique molecular compositions. This is supported by our observation of consistent SAP102 nanostructure at synapses without PSD-95 across development, while SAP102 at synapses with PSD-95 does change over time. For PSD- 95, our results regarding nano-organization during synapse maturation are generally consistent with those of Sun et al. (2022) who analyzed a partially overlapping developmental window and observed a similarly age-dependent increase in PSD-95 heterogeneity.

Another second intriguing possibility is that synapses with different compositions of MAGUKs may represent synapse classes with distinct presynaptic partners. There is evidence that synapses with PSD-95 only are associated with VGluT2 positive active zones while synapses with both proteins tend to be associated with VGluT1, suggesting that different compositions of MAGUKs are indicative of synapses of distinct origins (Zhu et al., 2018). Furthermore, while central elements of PSD-95 nanostructure are conserved across excitatory synapses onto pyramidal and parvalbumin-expressing interneurons, there are systematic, quantitative differences in its nanoscale distribution in that depend on the identity of the postsynaptic neuron (Dharmasri et al., 2023). We did not distinguish subsets of glutamatergic afferent neurons in these experiments, and thus paired or single-MAGUK synapse classes documented here may represent contacts formed between three neuron pair types, with accordingly unique rules of nanostructure architecture driving particular aspects of synapse function.

A third possibility for the mutual influence on nanostructure is that the presence of either SAP102 or PSD-95 impacts the nano-organization of the other either through shared interacting proteins, LLPS into common condensates, or perhaps even direct interactions with each other (Bonnet et al., 2013). PSD-95 dimerizes (Frank et al., 2017), and there is some evidence that SAP102 and PSD-95 may directly bind (Bonnet et al., 2013); however, while PSD-95 can form dimers and heterodimers with PSD-93, it apparently cannot do so with SAP102 (Zeng et al., 2018). Furthermore, it is possible that the MAGUKs compete for interactors, which may accentuate segregation. It is additionally intriguing that independent of whether SAP102 or PSD- 95 are alone or paired at synapses, they tend to have a similar number of NCs per synapse as each other, despite occupying a different fraction of the synapse as a whole. This suggests that other factors may be involved in establishing individual scaffold protein nanoclusters. An attractive model is that presynaptic release machinery via cell-adhesion molecules seed sites of MAGUK nanoclusters, which then in turn cluster other relevant synaptic proteins to maintain and modify synaptic function. Further multiplexed imaging using DNA-PAINT or other emerging technologies (Klevanski et al., 2020) to measure the receptors, signaling molecules, and other structural elements abundant within either paired or single-MAGUK NCs should help determine the role of distinct subsynaptic domains for maintaining and modifying synaptic function.

## Acknowledgements

This work was supported by F31 MH117920 and T32 GM008181 to P.A.D., F32 MH119687 to A.D.L, F31 MH124283 to M.C.A, and R37 MH080046 to T.A.B.

The authors thank Minerva Contreras for outstanding technical support, and the Blanpied lab for detailed discussions and evaluation of the manuscript.

The authors declare no competing financial interests.

---

**Figure 1-1.**
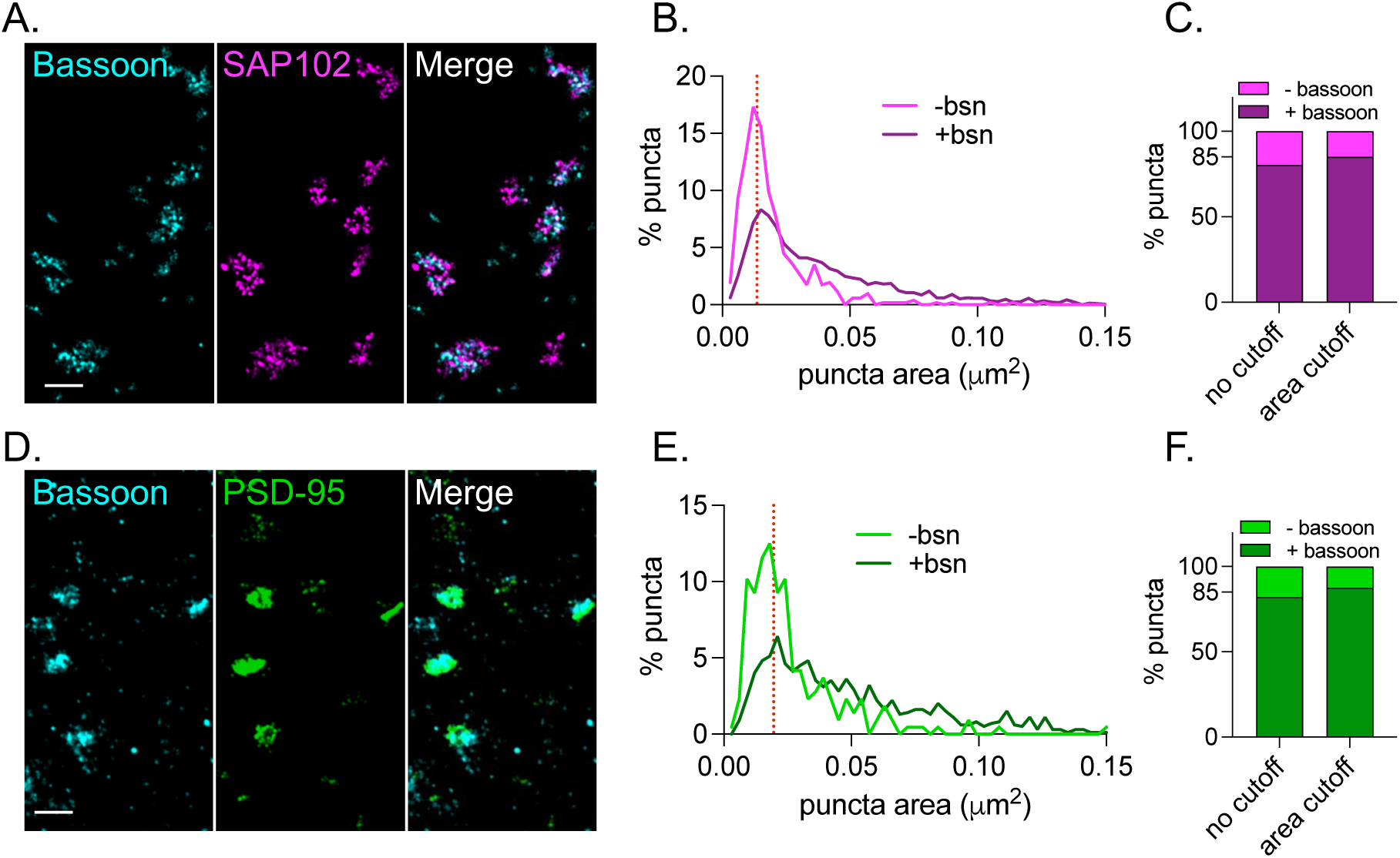
A. Example super-resolved synapses demonstrating co-localization between bassoon (left) and SAP102 (middle). B. Distribution of overall SAP102 puncta area at puncta that co-localized with bassoon and those that did not. Dashed line indicated cutoff value. C. Percentage of SAP102 puncta that overlapped with bassoon without the area cutoff applied and with the area cutoff applied. D. Example super-resolved synapses demonstrating co-localization between bassoon (left) and PSD-95 (middle). E. Distribution of overall PSD-95 puncta area at puncta that co-localized with bassoon and those that did not. Dashed line indicated cutoff value. F. Percentage of PSD-95 puncta that overlapped with bassoon without the area cutoff applied and with the area cutoff applied.

**Table 2-1:** Statistical analysis for Figure 2. Statistics related to data shown in Figure 2.

| FIG. 2A: SYNAPTIC PUNCTA AREA |  |  |  |
| --- | --- | --- | --- |
| DESCRIPTIVE STATISTICS |  |  |  |
| PROTEIN AND CATEGORY | Mean (mm<sup>2</sup>) | Standard deviation (mm<sup>2</sup>) | N (synapses) |
| SAP102 PAIRED | 0.05501 | 0.03413 | 602 |
| SAP102 ALONE | 0.02896 | 0.01725 | 455 |
| PSD-95 PAIRED | 0.09096 | 0.05207 | 602 |
| PSD-95 ALONE | 0.05378 | 0.03585 | 691 |
| TWO-WAY ANOVA/TUKEY'S MULTIPLE COMPARISON TEST |  |  |  |
| PROTEIN AND CATEGORY | Protein and category |  | p-value |
| SAP102 PAIRED | PSD-95 paired |  | <0.0001 |
|  | SAP102 alone |  | <0.0001 |
|  | PSD-95 alone |  | 0.9370 |
| PSD-95 PAIRED | SAP102 alone |  | <0.0001 |
|  | PSD-95 alone |  | <0.0001 |
| SAP102 ALONE | PSD-95 alone |  | <0.0001 |

| FIG. 2B: SYNAPTIC PUNCTA DENSITY |  |  |  |
| --- | --- | --- | --- |
| DESCRIPTIVE STATISTICS |  |  |  |
| PROTEIN AND CATEGORY | Mean (locs/mm<sup>2</sup>) | Standard deviation (locs/mm<sup>2</sup>) | N (synapses) |
| SAP102 PAIRED | 0.008714 | 0.003131 | 602 |
| SAP102 ALONE | 0.009175 | 0.006074 | 455 |
| PSD-95 PAIRED | 0.00901 | 0.003807 | 602 |
| PSD-95 ALONE | 0.006155 | 0.002874 | 691 |
| TWO-WAY ANOVA/TUKEY'S MULTIPLE COMPARISON TEST |  |  |  |
| PROTEIN AND CATEGORY | Protein and category |  | p-value |
| SAP102 PAIRED | PSD-95 paired |  | 0.7307 |
|  | SAP102 alone |  | 0.3188 |
|  | PSD-95 alone |  | <0.0001 |
| PSD-95 PAIRED | SAP102 alone |  | 0.9851 |
|  | PSD-95 alone |  | <0.0001 |
| SAP102 ALONE | PSD-95 alone |  | <0.0001 |

| FIG. 2E: NUMBER OF NCS PER SYNAPSE |  |  |  |
| --- | --- | --- | --- |
| DESCRIPTIVE STATISTICS |  |  |  |
| PROTEIN AND CATEGORY | Mean | Standard deviation | N (synapses) |
| SAP102 PAIRED | 2.126 | 1.322 | 602 |
| SAP102 ALONE | 1.499 | 0.8701 | 455 |
| PSD-95 PAIRED | 2.296 | 1.23 | 602 |
| PSD-95 ALONE | 1.437 | 0.8094 | 691 |

**TWO-WAY ANOVA/TUKEY'S MULTIPLE COMPARISON TEST**
| PROTEIN AND CATEGORY | Protein and category | p-value |
| --- | --- | --- |
| SAP102 PAIRED | PSD-95 paired | 0.0331 |
|  | SAP102 alone | <0.0001 |
|  | PSD-95 alone | <0.0001 |
| PSD-95 PAIRED | SAP102 alone | <0.0001 |
|  | PSD-95 alone | <0.0001 |
| SAP102 ALONE | PSD-95 alone | 0.7791 |

**DESCRIPTIVE STATISTICS**
| PROTEIN AND CATEGORY | Mean (nm<sup>2</sup>) | Standard deviation (nm<sup>2</sup>) | N (NCs) |
| --- | --- | --- | --- |
| SAP102 PAIRED | 2206 | 3245 | 1280 |
| SAP102 ALONE | 1478 | 1505 | 682 |
| PSD-95 PAIRED | 3609 | 4487 | 1382 |
| PSD-95 ALONE | 2624 | 2713 | 993 |

**TWO-WAY ANOVA/TUKEY'S MULTIPLE COMPARISON TEST**
| PROTEIN AND CATEGORY | Protein and category | p-value |
| --- | --- | --- |
| SAP102 PAIRED | PSD-95 paired | <0.0001 |
|  | SAP102 alone | <0.0001 |
|  | PSD-95 alone | 0.0192 |
| PSD-95 PAIRED | SAP102 alone | <0.0001 |
|  | PSD-95 alone | <0.0001 |
| SAP102 ALONE | PSD-95 alone | <0.0001 |

**DESCRIPTIVE STATISTICS**
| PROTEIN AND CATEGORY | Mean (locs/nm<sup>2</sup>) | Standard deviation (locs/nm<sup>2</sup>) | N (synapses) |
| --- | --- | --- | --- |
| SAP102 PAIRED | 0.08289 | 0.06884 | 602 |
| SAP102 ALONE | 0.07932 | 0.05704 | 455 |
| PSD-95 PAIRED | 0.08909 | 0.05271 | 602 |
| PSD-95 ALONE | 0.07286 | 0.04787 | 691 |

**TWO-WAY ANOVA/TUKEY'S MULTIPLE COMPARISON TEST**
| PROTEIN AND CATEGORY | Protein and category | p-value |
| --- | --- | --- |
| SAP102 PAIRED | PSD-95 paired | 0.2313 |
|  | SAP102 alone | 0.7428 |
|  | PSD-95 alone | 0.0085 |
| PSD-95 PAIRED | SAP102 alone | 0.0291 |
|  | PSD-95 alone | <0.0001 |
| SAP102 ALONE | PSD-95 alone | 0.2356 |

| FIG. 2J: NC DENSITY |  |  |  |
| --- | --- | --- | --- |
| DESCRIPTIVE STATISTICS |  |  |  |
| PROTEIN AND CATEGORY | Mean (locs/nm<sup>2</sup>) | Standard deviation (locs/nm<sup>2</sup>) | N (NCs) |
| SAP102 PAIRED | 0.05562 | 0.0434 | 1280 |
| SAP102 ALONE | 0.0635 | 0.05681 | 682 |
| PSD-95 PAIRED | 0.04735 | 0.02989 | 1382 |
| PSD-95 ALONE | 0.03384 | 0.0222 | 993 |
| TWO-WAY ANOVA/TUKEY'S MULTIPLE COMPARISON TEST |  |  |  |
| PROTEIN AND CATEGORY | Protein and category | p-value |  |
| SAP102 PAIRED | PSD-95 paired | <0.0001 |  |
|  | SAP102 alone | <0.0001 |  |
|  | PSD-95 alone | <0.0001 |  |
| PSD-95 PAIRED | SAP102 alone | <0.0001 |  |
|  | PSD-95 alone | <0.0001 |  |
| SAP102 ALONE | PSD-95 alone | <0.0001 |  |

| FIG. 2K: NC DENSITY |  |  |  |
| --- | --- | --- | --- |
| DESCRIPTIVE STATISTICS |  |  |  |
| PROTEIN AND CATEGORY | Mean (locs/nm<sup>2</sup>) | Standard deviation (locs/nm<sup>2</sup>) | N (synapses) |
| SAP102 PAIRED | 0.02763 | 0.02207 | 602 |
| SAP102 ALONE | 0.04089 | 0.04289 | 455 |
| PSD-95 PAIRED | 0.02537 | 0.01434 | 602 |
| PSD-95 ALONE | 0.02072 | 0.0134 | 691 |
| TWO-WAY ANOVA/TUKEY'S MULTIPLE COMPARISON TEST |  |  |  |
| PROTEIN AND CATEGORY | Protein and category | p-value |  |
| SAP102 PAIRED | PSD-95 paired | 0.3678 |  |
|  | SAP102 alone | <0.0001 |  |
|  | PSD-95 alone | <0.0001 |  |
| PSD-95 PAIRED | SAP102 alone | <0.0001 |  |
|  | PSD-95 alone | 0.0033 |  |
| SAP102 ALONE | PSD-95 alone | <0.0001 |  |

**Table 4-1:**
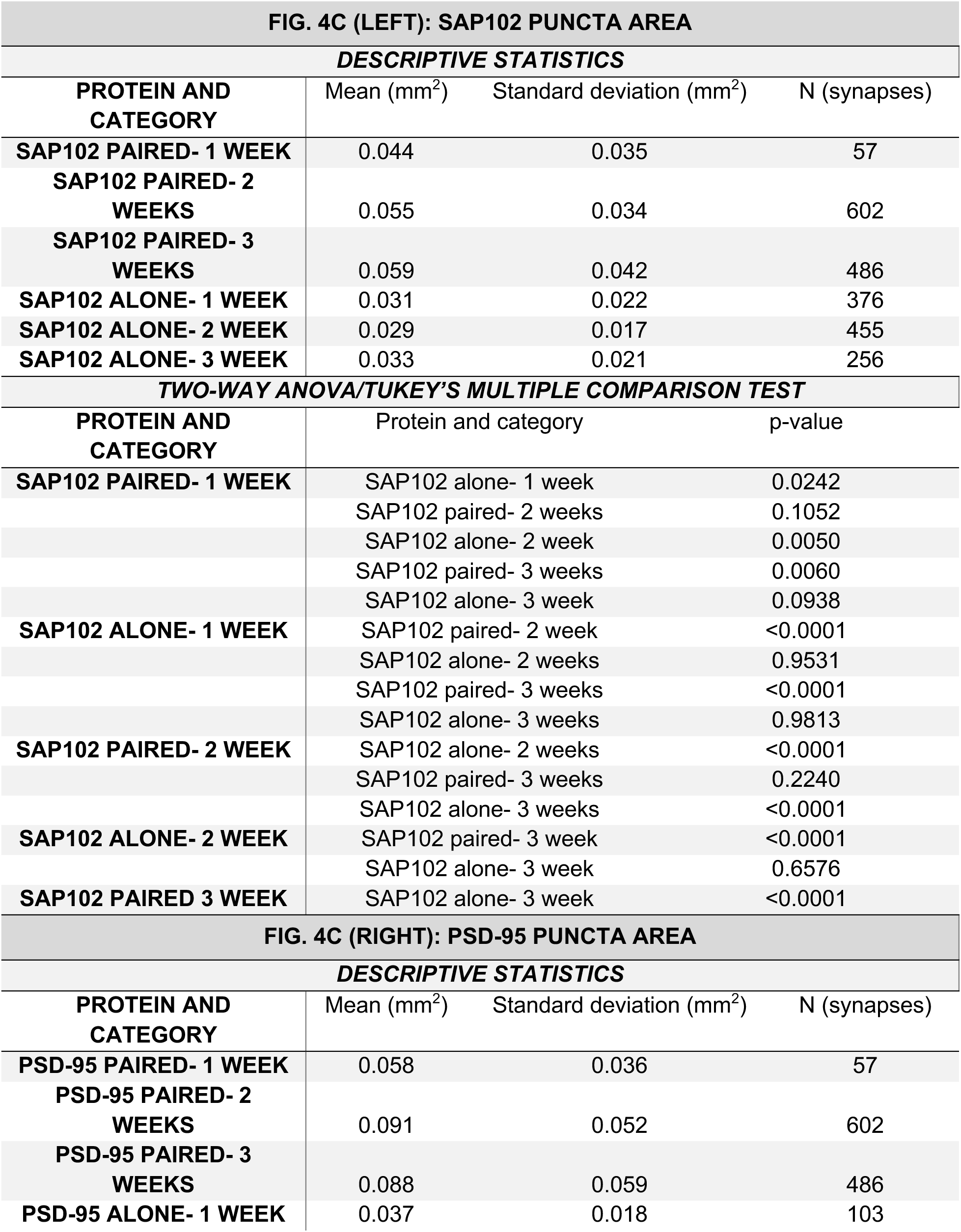
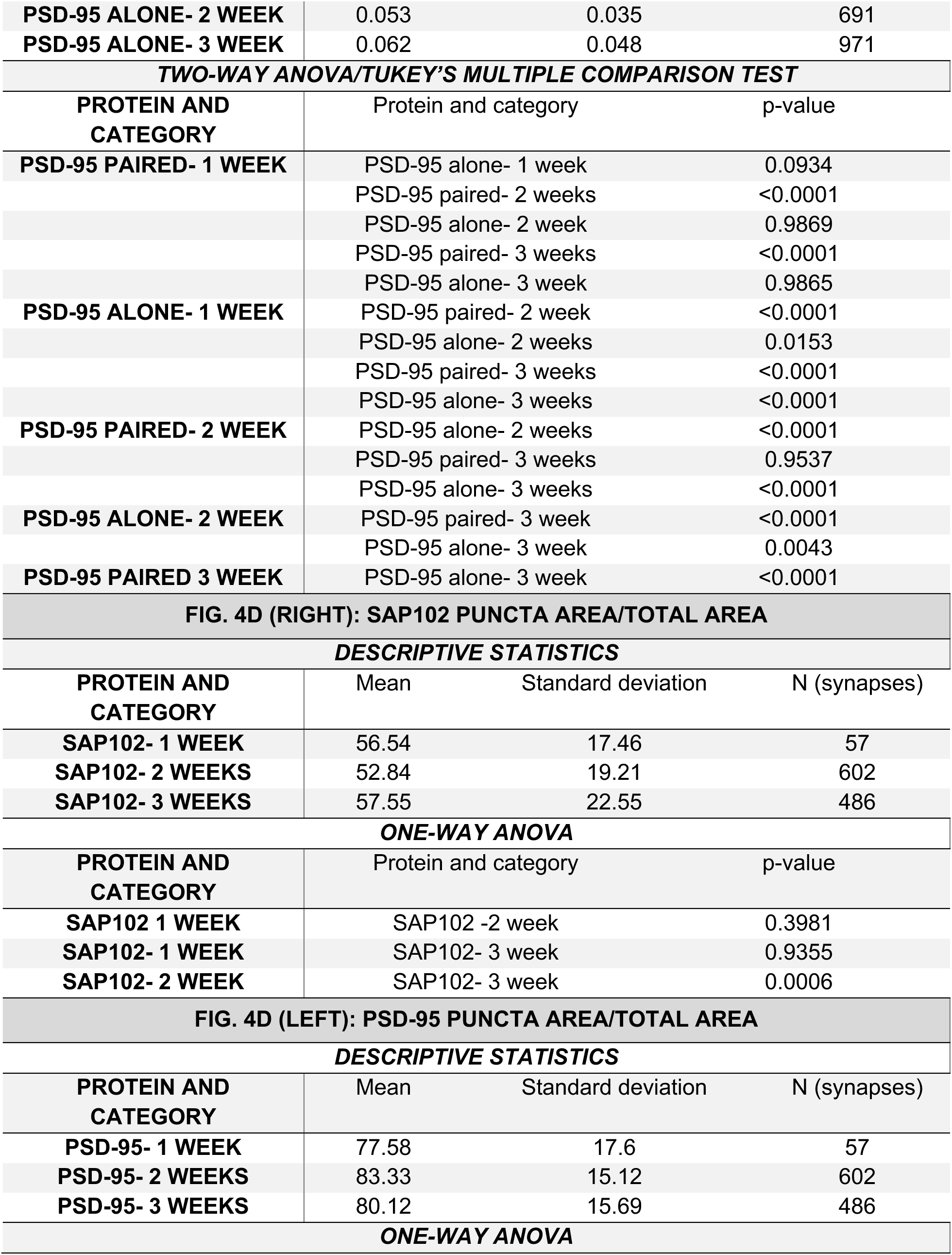
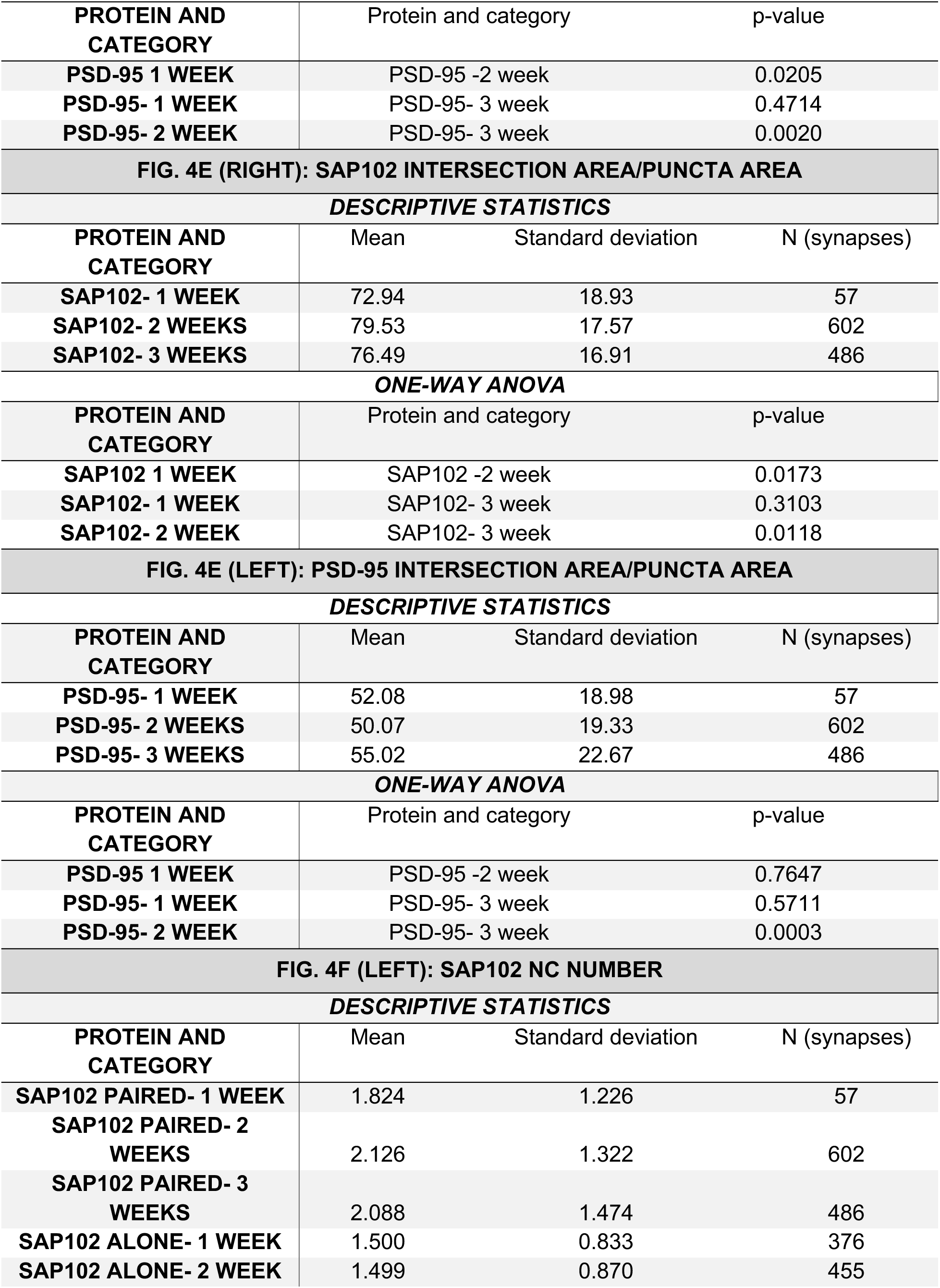
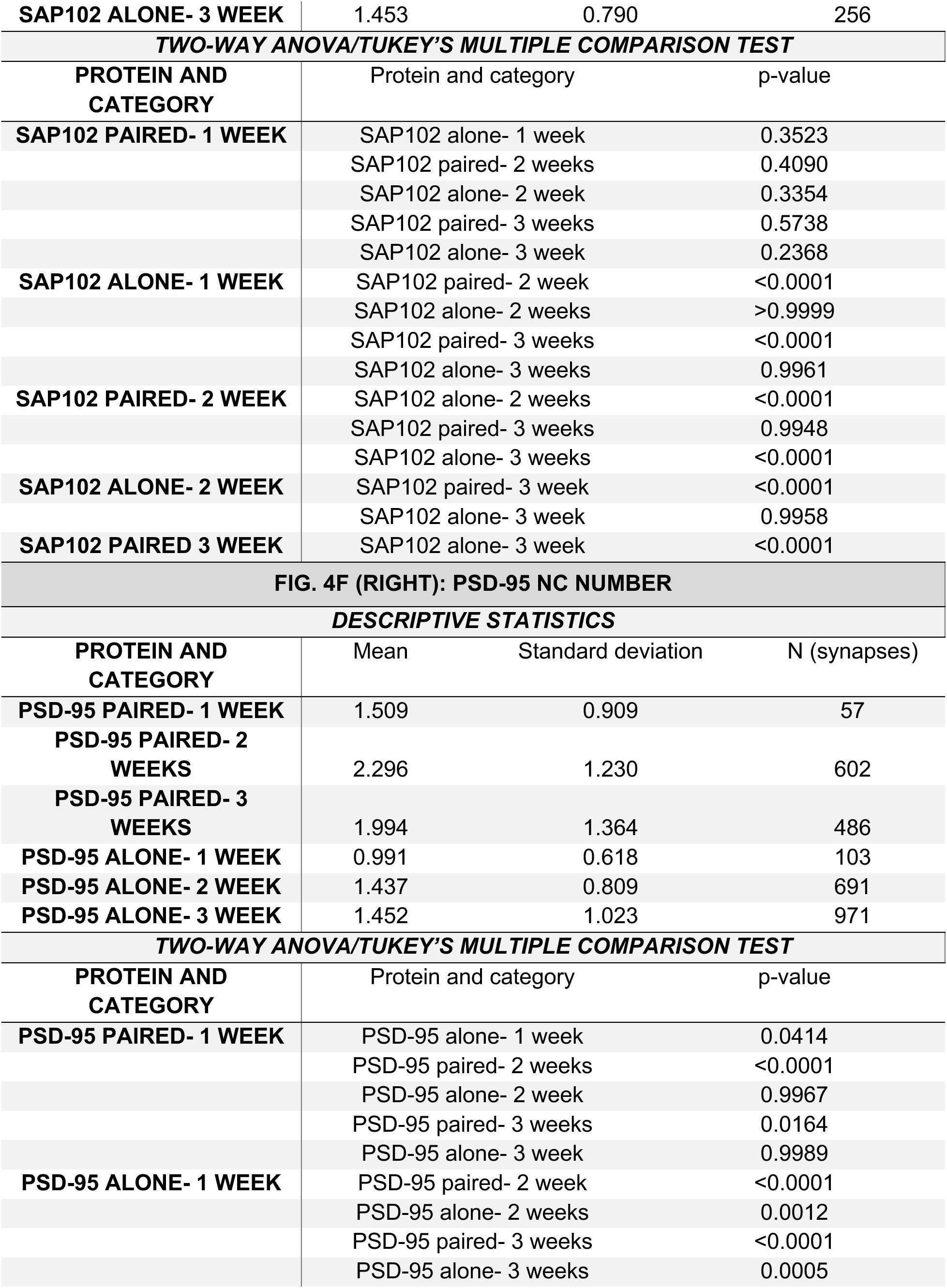
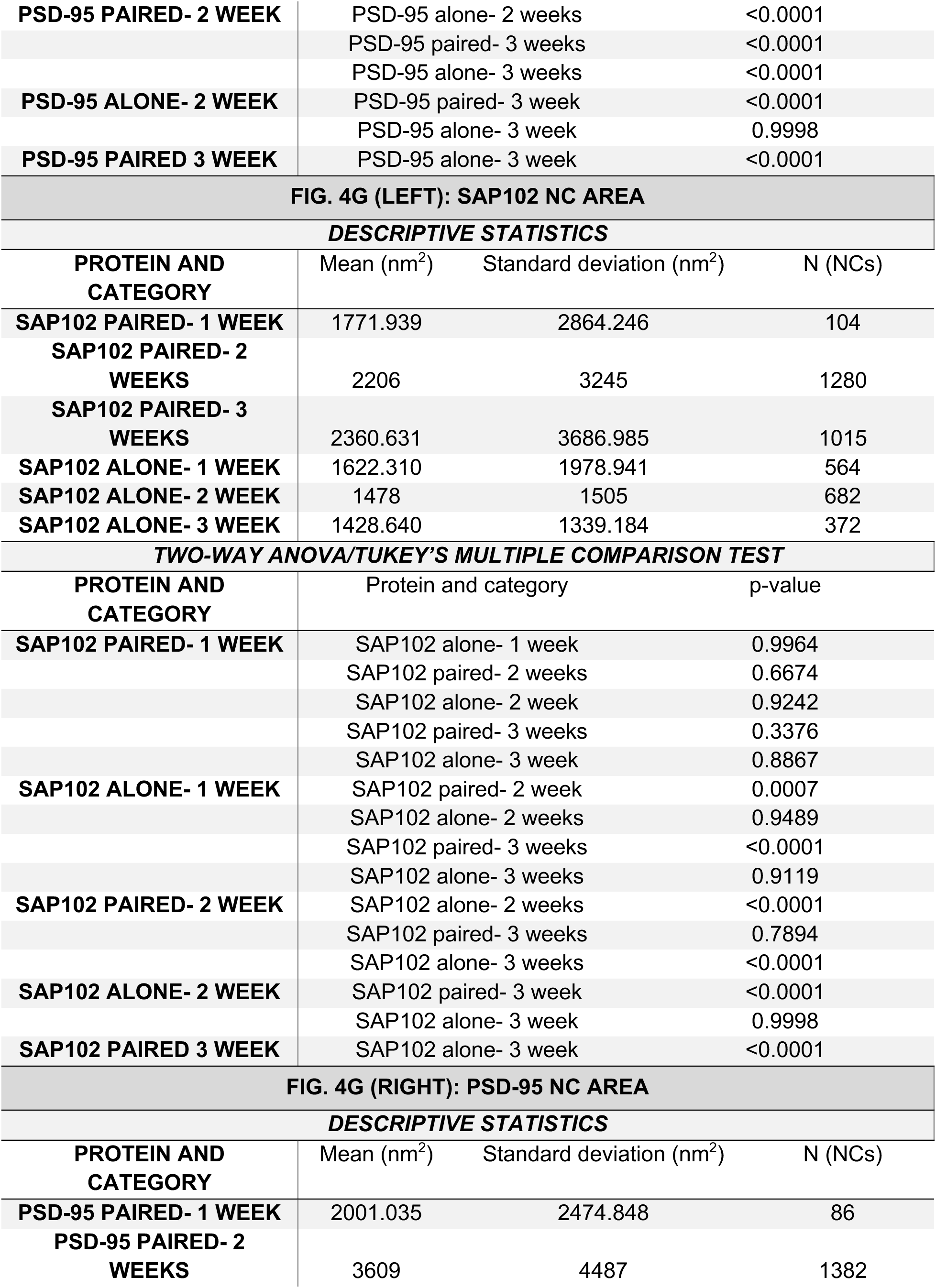
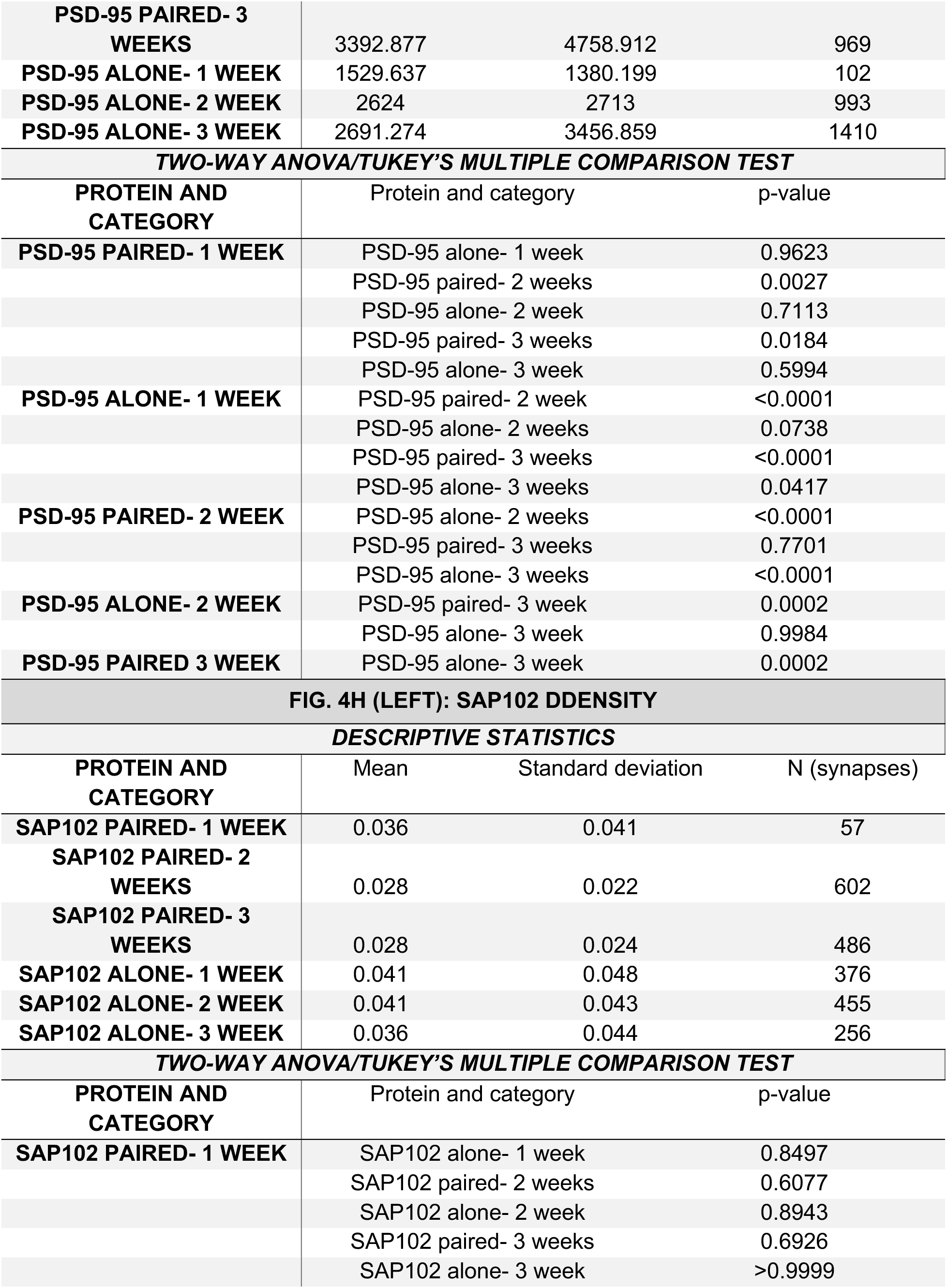
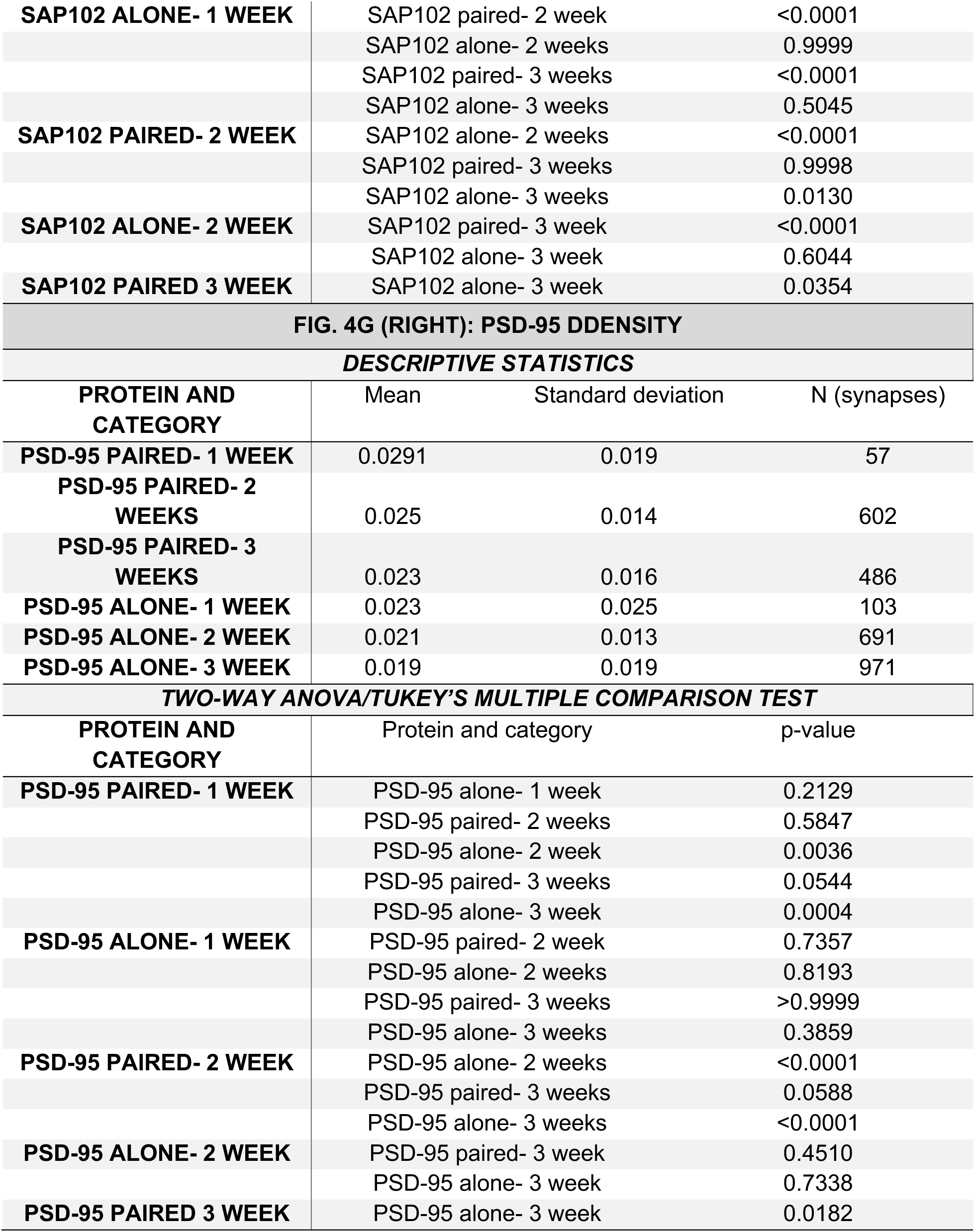
Statistical analysis for Figure 4. Statistics related to data shown in Figure 4.

## Notes

### Competing Interest Statement

The authors have declared no competing interest.

